# Methyl-Metabolite Depletion Elicits Adaptive Responses to Support Heterochromatin Stability and Epigenetic Persistence

**DOI:** 10.1101/726448

**Authors:** Spencer A. Haws, Deyang Yu, Cunqi Ye, Coral K. Wille, Long C. Nguyen, Kimberly A. Krautkramer, Jay L. Tomasiewicz, Shany E. Yang, Blake R. Miller, Wallace H. Liu, Kazuhiko Igarashi, Rupa Sridharan, Benjamin P. Tu, Vincent L. Cryns, Dudley W. Lamming, John M. Denu

**Author notes:** Correspondence: John M. Denu.

## Abstract

S-adenosylmethionine (SAM) is the methyl-donor substrate for DNA and histone methyltransferases that regulate cellular epigenetic states. This metabolism-epigenome link enables the sensitization of chromatin methylation to altered SAM abundance. However, a chromatin-wide understanding of the adaptive/responsive mechanisms that allow cells to actively protect epigenetic information during life-experienced fluctuations in SAM availability are unknown. We identified a robust response to SAM depletion that is highlighted by preferential cytoplasmic and nuclear *de novo* mono-methylation of H3 Lys 9 (H3K9) at the expense of global losses in histone di- and tri-methylation. Under SAM-depleted conditions, *de novo* H3K9 mono-methylation preserves heterochromatin stability and supports global epigenetic persistence upon metabolic recovery. This unique chromatin response was robust across the mouse lifespan and correlated with improved metabolic health, supporting a significant role for epigenetic adaptation to SAM depletion *in vivo*. Together, these studies provide the first evidence for active epigenetic adaptation and persistence to metabolic stress.

## Introduction

Metabolism and the epigenome are connected by central metabolites that act as co-substrates for chromatin modifying enzymes, enabling fluctuations in metabolite availability to directly tune a cell’s ability to ‘write’ and ‘erase’ chromatin post-translational modifications (PTMs) (Fan et al., 2015; Li et al., 2018). Chromatin methylation has been shown to be particularly susceptible to metabolic perturbations as methyltransferases require a single methyl-donor cofactor, S-adenosylmethionine (SAM) (Ducker and Rabinowitz, 2016; Mentch and Locasale, 2017). For example, accumulation of SAM in *S. cerevisiae* results in global increases in histone methylation levels (Ye et al., 2017). On the contrary, reduced SAM availability has been linked with site-specific losses in histone methylation as well as global depression of DNA methylation in several organisms (Shiraki et al., 2014; Ding et al., 2015; Kera et al., 2013; Tang et al., 2017; Shyh-Chang et al., 2013; Mentch et al., 2015; Tang et al., 2015; Hayashi et al., 2018; Towbin et al., 2012; Strekalova et al., 2019; Wang et al., 2019).

Intracellular SAM abundance is dependent on the catalysis of its obligatory precursor metabolite, the essential amino acid methionine (Met). Consumption of popular diets containing low quantities of Met (fish, vegetarian, and vegan) have been shown to significantly correlate with decreased plasma Met concentrations (Farmer, 2014; Schmidt et al., 2016). This link provides an opportunity for diet to directly influence intracellular SAM availability. Furthermore, abundance of Met and SAM have both been shown to fluctuate naturally in tune with circadian rhythms (Krishnaiah et al., 2017). Therefore, individuals consuming adequate amounts of dietary Met likely still experience periods of decreased methyl-metabolite availability. While both dietary intake and circadian regulation are capable of influencing Met and SAM abundance, adaptive mechanisms that allow cells to actively respond to – and recover their functional epigenomes from – such metabolic perturbations are unknown. Here, we define the ability of a cell and/or organism to re-establish the epigenome that was present prior to onset of an environmental perturbation or metabolic stress as *epigenetic persistence*.

Adaptive mechanisms that respond to methyl-metabolite depletion likely support critical cellular functions under these conditions. Interestingly, dietary Met-restriction is associated with beneficial metabolic reprogramming and lifespan extension in mammals (Orentreich et al., 1993; Yu et al., 2018; Miller et al., 2005; Brown-Borg et al., 2018; Lees et al., 2014; Green and Lamming, 2018). These phenotypes run counter to negative pathologies associated with dysregulated chromatin methylation (i.e., intellectual disability syndromes, cancers, and premature as well as natural aging) (Greer and Shi, 2012). This suggests cells possess responsive mechanisms capable of actively maintaining regulation of site specific and/or global chromatin methylation events during SAM depletion. The goal of this study is to identify and characterize such mechanisms to better understand how organisms actively protect their epigenome in response to fluctuations in central metabolite availability.

Here, we identify a robust, conserved chromatin response to metabolically-depleted SAM levels. This cellular response is characterized by global decreases in higher-state histone methylation (di- and tri-methyl) and simultaneous coordinated activation of *de novo* H3K9 mono-methylation involving newly-synthesized and chromatin-bound histone H3. Under extreme conditions of both reduced SAM levels and inhibited methylation that would yield H3K9me1, heterochromatin instability and de-repression of constitutively silenced DNA elements is exacerbated. Acute inhibition of *de novo* H3K9 mono-methylation also leads to the dysregulation of global histone PTM states upon metabolic recovery, disrupting epigenetic persistence to decreased SAM availability. Importantly, chronic metabolic depletion of SAM in both young and old mice recapitulates these *in vitro* findings, revealing a conserved, age-independent chromatin response. These results implicate *de novo* H3K9 mono-methylation as an indispensable mechanism to support heterochromatin stability and global epigenetic persistence in response to SAM depletion.

## Results

### Methionine Restriction Stimulates Dynamic Histone PTM Response

To comprehensively investigate how disruption of methyl-donor metabolism affects chromatin methylation states, global histone proteomics and DNA methylation (5mC) analyses were performed on two distinct *in vitro* and *in vivo* systems. These included Met-restricted HCT116 human colorectal cancer cells and C57BL/6J mouse liver. Methionine metabolism was severely repressed in both systems, highlighted by significant reductions in Met and SAM abundance (Figure 1A and Figure S1A-S1D). Reduced methyl-metabolite availability did not affect global DNA 5mC abundance in either system (Figure 1B-1C). However, LC-MS/MS analysis of more than 40 unique histone H3 peptide proteoforms revealed dynamic PTM responses to Met-restriction in both HCT116 cells and C57BL/6J liver (Figure 1D-1E). In HCT116 cells, Met-restriction stimulated distinct biphasic changes in global PTM profiles (Figure 1D). Phase I (0 hr-45 min) was rapid and marked by a trending upregulation of H3K4me2/3, PTMs known to mark transcriptionally active promoters. Phase II (90 min-24 hrs) was characterized by global decreases in di- and tri-histone methylation. Decreased levels of di- and tri-methylated peptides were accompanied by increases in acetylated and unmodified peptide species. Similarly in C57BL/6J mice, histone PTM responses were marked by significant decreases in histone di- and tri-methylation that essentially matched the patterns found in HCT116 cells during the prolonged Phase II response (Figure 1E and Figure S1E-S1J). Decreased higher state (di- and tri-) histone methylation both *in vitro* and *in vivo* highlights a decreased methylation capacity resulting from prolonged Met and/or SAM depletion. Together, these observations suggest perturbed methyl-donor metabolism is capable of stimulating significant changes in histone, but not global 5mC, methylation abundance. Furthermore, similarities between the metabolic and epigenetic responses to Met-restriction across both systems support the use of *in vitro* Met-restriction as a model for mechanistic follow-up studies.

**Figure 1.**
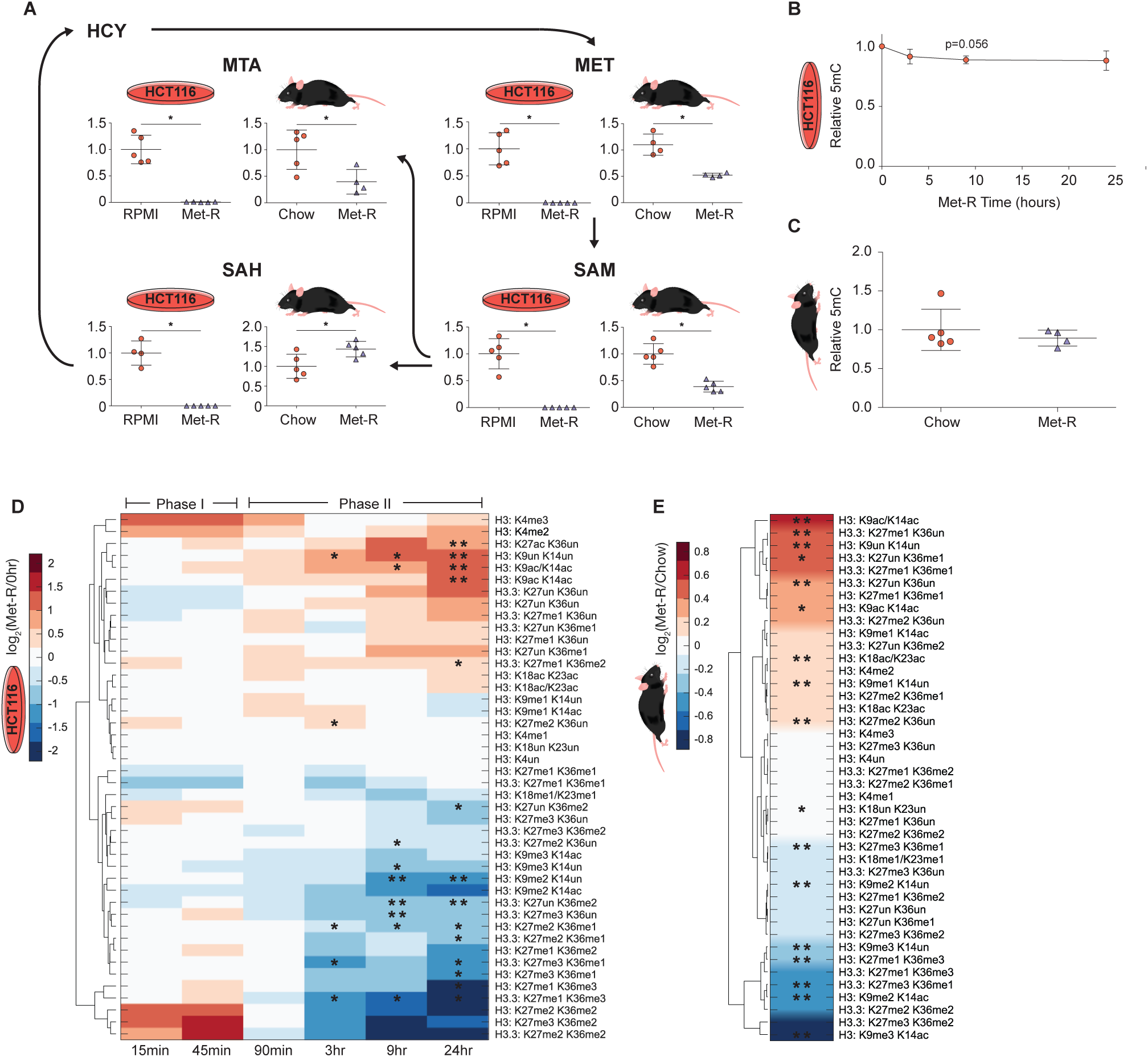
Methionine Restriction Stimulates Dynamic Histone PTM Response (A) Metabolic pathway diagram illustrating changes in Met-cycle metabolite abundance after 24 hours and 5 weeks of Met-restriction in HCT116 cells and C57BL/6J liver, respectively. n=5, error bars represent SD, *p<0.05 (Welch’s t-Test). (*B-C*) Plots illustrating global, relative 5mC DNA methylation levels during Met-restriction. n≥3, error bars represent SD, *p<0.05 (Welch’s t-Test) (*D-E*) Hierarchichal clustered heatmaps of LC-MS/MS generated log_2_ fold-change stoichiometric histone H3 peptide proteoform values relative to 0-hour or chow diet controls in HCT116 cells and C57BL/6J liver, respectively. n 3, *p<0.05, **p<.01 (Welch’s t-Test). See also Table S1 and Figure S1.

### Decreased SAM Availability Drives Robust Histone Methylation Response

Dramatic reduction of intracellular Met and SAM correlated with onset of the *in vitro* Phase I and II histone PTM changes, respectively (Figure 1D and Figure S1A-S1B). This implies depletion of individual methyl-metabolites may be capable of stimulating distinct histone modifying pathways. To determine if the global losses in di- and tri-histone methylation *in vitro* are driven specifically by SAM depletion, two alternative approaches were employed to deplete cellular SAM independent of Met (Figure 2A). The first approach utilized RNAi knockdown of the mammalian SAM synthetase MAT2A (methionine adenosyltransferase II alpha). Knockdown via MAT2A-RNAi treatment decreased MAT2A transcript abundance by greater than 97%, resulting in a near complete deprivation of SAM availability with no effect on Met levels (Figure 2B-2D). Histone proteomics analysis of MAT2A-RNAi treated cells identified site-specific histone PTM changes that mirrored those identified in HCT116 cells under Phase II Met-restriction conditions (r=0.778, p=8.59e_-10_) (Figure 2E). Using a complementary approach, SAM levels were depleted by overexpression of a major SAM consumer, PEMT (phosphatidylethanolamine N-methyltransferase). Ye et al. 2017 have previously shown PEMT overexpression significantly reduces SAM availability in HEK293T human embryonic kidney cells. Histone proteomics analysis of PEMT overexpressing HEK293T cells identified site-specific PTM changes that paralleled those of Met-restricted controls (r=0.803, p=3.30e_-8_) (Figure 2F). Together, two separate means of reducing SAM levels demonstrate SAM reduction alone is sufficient to inhibit the methylation capacity of a cell, inducing dynamic changes in histone PTM abundance. Furthermore, these findings support the use of prolonged Met-restriction as a model of SAM depletion.

**Figure 2.**
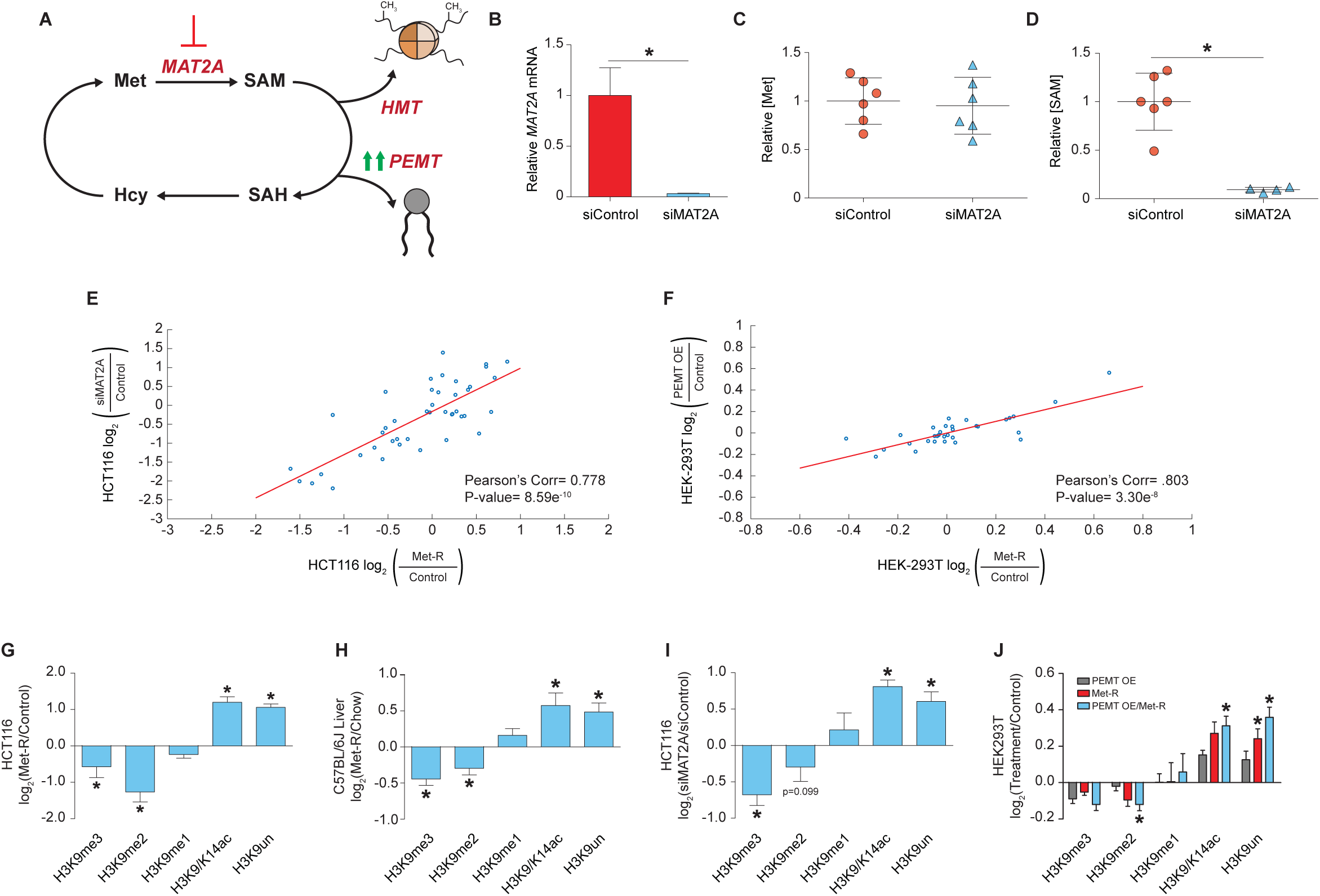
Decreased SAM Availability Drives Robust Histone Methylation Response (*A*) Pathway diagram illustrating experimental approaches to depleting intracellular SAM availability. (*B*) Bar graph of relative *MAT2A* mRNA abundance in HCT116 cells as measured Met and SAM levels in HCT116 cells. n 4, *p<0.05 (Welch’s t-test). (*E-F*) Correlation plot of LC- MS/MS generated log_2_ fold-change stoichiometric values for individual histone H3 peptide proteoforms. n=3. (*G-*J) Bar graphs illustrating LC-MS/MS generated log_2_ fold-changes for individual H3K9 PTMs. n=3 *p<0.05 (Welch’s t-test) See also Table S1, Table S2, and Figure S2.

The similar changes in global histone PTM abundance across four SAM depletion systems were analyzed for the presence of methyl PTMs whose abundances either increased or were unaffected by this metabolic perturbation. Such PTMs would be candidate mediators of an adaptive response to SAM depletion. Unexpectedly, all four SAM depletion systems exhibited identical changes in H3K9 methylation (Figure 2G-2J). SAM depletion stimulated a decrease in H3K9me2/3 with a corresponding increase in H3K9ac/14ac and unmodified peptides while net H3K9me1 abundance was unchanged. Preservation of net H3K9me1 levels in response to SAM depletion was reproducible in Met-restricted MCF7, HEK-293, Panc1, and HepG2 cell lines, as well as in MAT2A-RNAi treated HepA1 cells (Figure S2A-S2F). The robust conservation of H3K9me1 abundance in diverse systems marked by reduced methylation capacity suggest this PTM may be supported by adaptive, active mechanisms.

### H3K9me1 Abundance under SAM Depletion is Driven by *de novo* Methylation

Global levels of H3K9me1 could be maintained through three distinct mechanisms during SAM depletion: 1.) protection from PTM turnover, 2.) de-methylation of H3K9me2/3, and/or 3.) *de novo* methylation of unmodified H3K9. The decrease in H3K9me2/3 and increase in unmodified H3K9 peptide levels across SAM depletion systems suggest H3K9 can be subjected to facile demethylation while *de novo* methylation appears enzymatically unfavorable. However, *de novo* H3K9 mono-methylation would constitute an active, adaptive response to this metabolic stress that is capable of supporting critical chromatin functions.

To investigate the contribution of *de novo* methylation to global H3K9me1 levels, RNAi-mediated repression of five H3K9 HMTs was first performed in Met-replete conditions to determine which enzymes catalyzed a majority of H3K9 mono-methylation reactions in HCT116 cells. Repression of HMT expression had significant yet restricted effects, given technical limitations, on H3K9 PTM abundance and suggested Ehmt1/Ehmt2 were the primary H3K9 mono-methyltransferases in this system (Figure S3A-S3D). Because repression of HMT expression was intended to be coupled with Met-restriction, RNAi treatment could not be extended to elicit larger changes in PTM abundance as HCT116 cells no longer proliferate beyond 24 hours of Met-restriction (Figure S3E). Therefore, small molecule (UNC0642) inhibition of H3K9 HMTs Ehmt1 and Ehmt2 was used as the best approach to acutely block nuclear H3K9 mono-methylation (Liu F et al., 2013). UNC0642 treatment of Met-replete cells reduced H3K9me1 abundance to a greater extent than RNAi repression of EHMT1/2 (Figure 3A and Figure S3D). Coupling UNC0642 treatment with Met-restriction also stimulated a significant decrease in H3K9me1 levels (Figure 3B and Figure S3F). Comparatively, these data suggest Ehmt1/Ehmt2 retain nearly 60% of their H3K9 mono-methylation activity during severe *in situ* SAM deprivation and that this activity is critical for preserving global H3K9me1 abundance (Figure 3C). Furthermore, H3K9me2 abundance decreased significantly in Met-replete cells upon UNC0642 treatment but was unaffected when UNC0642 was coupled with Met-restriction (Figure 3A-3B). This indicates only the mono-methylation activity of Ehmt1 and Ehmt2 is preferentially supported during SAM depletion.

**Figure 3.**
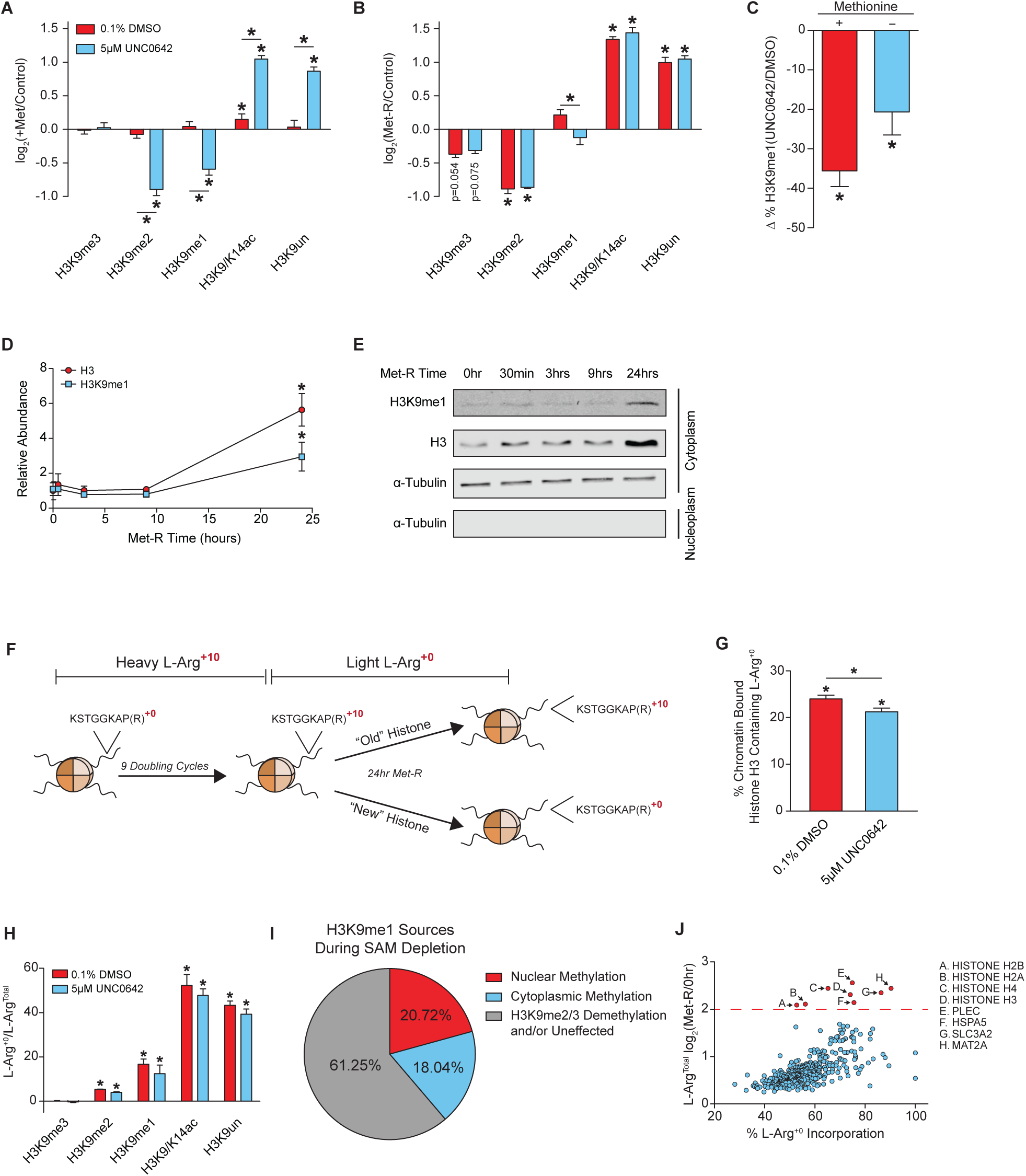
H3K9me1 Abundance under SAM Depletion is Driven by *de novo* Methylation in HCT116 cells. n≥, error bars represent SD, *p<0.05 (Welch’s t-Test). (*C*) Bar graph UNC0642 treatment. n≥2, error bars represent SD, *p<0.05 (Welch’s t-Test) (*D*) Scatter plot of illustrating percent loss in global H3K9me1, as measured by LC-MS/MS, in response to 5*μ*m relative, total H3 and H3K9me1 cytoplasmic protein abundance in HCT116 cells as measured by western blot. Error bars represent SD, n=3, *p<0.05 (Welch’s t-Test). (*E*) Representative western blot images of those used for the quantification presented in Figure 3E. Alpha-Tubulin is used as a cytoplasm marker to confirm efficient sub-cellular fractionation. See Document S1 for REVERT Total Protein Stain images used for normalization. (*F*) Diagram illustrating the SILAC experimental design. (*G*) Bar graph illustrating contribution of newly synthesized histone H3 to the total chromatin bound pool after 24 hours of Met-restriction in HCT116 cells as measured by LC-MS/MS. n 2, error bars represent SD, *p<0.05 (Welch’s t-test). (*H*) Bar graph illustrating ratio of light:heavy L-Arg isotope presence in H3K9 proteoforms after 24 hours of 2, Error bars represent SD, *p<0.05 (Welch’s t-Test). (*I*) Pie chart depicting sources of H3K9me1 during SAM depletion. (*J*) Scatter plot of 373 cytoplasmic proteins identified by LC-MS/MS with significant incorporation of L-Arg _+0_ that were also significantly upregulated after 24 hours of Met-restriction in 0.1% DMSO treated HCT116 cells. n=4, Statistical significance of p<0.05 (Welch’s t-Test). See also Table S1, Table S3, and Figure S3.

In addition to methylation of H3K9 by nuclear enzymes, mono-methylation is known to be enzymatically catalyzed on cytoplasmic histones prior to nuclear import and deposition onto chromatin (Loyola A et al., 2006; Pinheiro et al., 2012). Subcellular fractionation of Met-restricted HCT116 cells coupled with western blot analyses indicated that cytoplasmic H3K9me1 levels increase 3-fold after 24 hours of Met-restriction, with a similar increase in total H3 protein levels (Figure 3D-3E). To determine the contribution of cytoplasmic H3K9me1 to the chromatin-bound nuclear pool after 24 hours of Met-restriction, a pulse-chase, SILAC experiment was performed (Figure 3F). L-Arg_+10_ isotope incorporation prior to Met-restriction was determined to be nearly complete with average incorporation for H3K9 peptides greater than 98% (Figure S3H). Newly synthesized H3 was calculated to comprise 24% of the chromatin bound histone pool and contributed 18% of the global H3K9me1 signal after 24 hours of Met-restriction (Figure 3G-3H). UNC0642 treatment did not significantly lessen the contribution of cytoplasmic H3K9me1 to the total pool, suggesting cytoplasmic H3K9me1 may be largely protected from enzyme catalyzed turnover once deposited onto chromatin under these conditions (Figure 3H). Modest, insignificant decreases in L-Arg_+0_ H3K9 PTM signal after UNC0642 treatment can be attributed to reduced incorporation of newly synthesized H3 onto chromatin. (Figure 3G) Reduced H3 incorporation in UNC0642-treated cells is likely the consequence of a slower growth rate under these conditions (Figure S3E). Together, these results suggest ∼40% of global H3K9me1 abundance is driven by *de novo* methylation during Met-restriction with nearly equal contributions from both the cytoplasm and nucleus. The remaining ∼60% of H3K9me1 levels are likely protected from turnover and/or are generated via H3K9me2/3 de-methylation during SAM depletion (Figure 3I).

The significant upregulation of cytoplasmic H3K9me1 and total H3, coupled with their incorporation onto chromatin during SAM depletion, were unexpected. These findings suggest cells utilize scarcely available Met and SAM to support the protein abundance and methylation of histone H3 under these conditions. Shotgun proteomics was performed on cytoplasmic protein fractions of the previously described SILAC experiment to determine if histone protein abundance is uniquely responsive to SAM depletion. Of the more than 2,400 identified proteins, only 8 possessed significant incorporation of the light L-Arg_+0_ isotope and also increased in absolute abundance by a log_2_ fold-change value greater than 2.0 (Figure 3J). Frequency of Met residues in the primary amino acid sequence did not affect L-Arg_+0_ isotope incorporation (Figure S3I). Remarkably, all four core histone subunits, Mat2A, and the methionine importer SLC3A2 (Bröer et al., 2001) were among the 8 proteins (Figure 3J). These results suggest proteins critical for chromatin stability and methyl-donor metabolism are preferentially generated during metabolically-induced SAM depletion. To our knowledge, this is also the first example of a metabolic stress stimulating increased histone protein abundance.

### H3K9me1 is Redistributed Over Repetitive and Transposable Elements During SAM Depletion

Multiple, non-redundant mechanisms for H3K9me1 maintenance suggest strong biological pressure exists to retain this PTM under SAM depleted conditions. One critical function for H3K9me1 is to serve as a primer for H3K9me2/3 methylation as methyltransferases that yield these higher methylation states require a mono-methyl substrate (Peters et al., 2001; Loyola et al., 2006; Loyola et al., 2009). Accordingly, installment of H3K9me2/3 for constitutive heterochromatin-mediated repression of telomeric, pericentromeric, and centromeric repetitive/transposable elements requires readily available H3K9me1 (Rea et al., 2000; Bannister et al., 2001; Lachner et al., 2000).

To determine if H3K9me1 is being maintained globally or redistributed to regions of susceptible heterochromatin during SAM depletion, ChIP-sequencing was performed. Met-restriction stimulated a 35**%** loss in total H3K9me3 peak abundance (Figure 4A). A higher percentage of H3K9me3 peaks were lost at LTR, intergenic, and intron annotated loci relative to the total, while H3K9me3 enrichment at satellite, simple repeat, and SINE annotated loci were slightly less sensitive to SAM depletion (Figure 4A). Interestingly, H3K9me1 enrichment at repetitive and transposable loci were either largely unaffected by SAM depletion or increased (Figure 4B). Only the percent change for intergenic and intron annotated H3K9me1 peaks was lost to a greater extent than the total peak number, suggesting H3K9me1 is more susceptible to de-methylation at these genomic loci.

**Figure 4.**
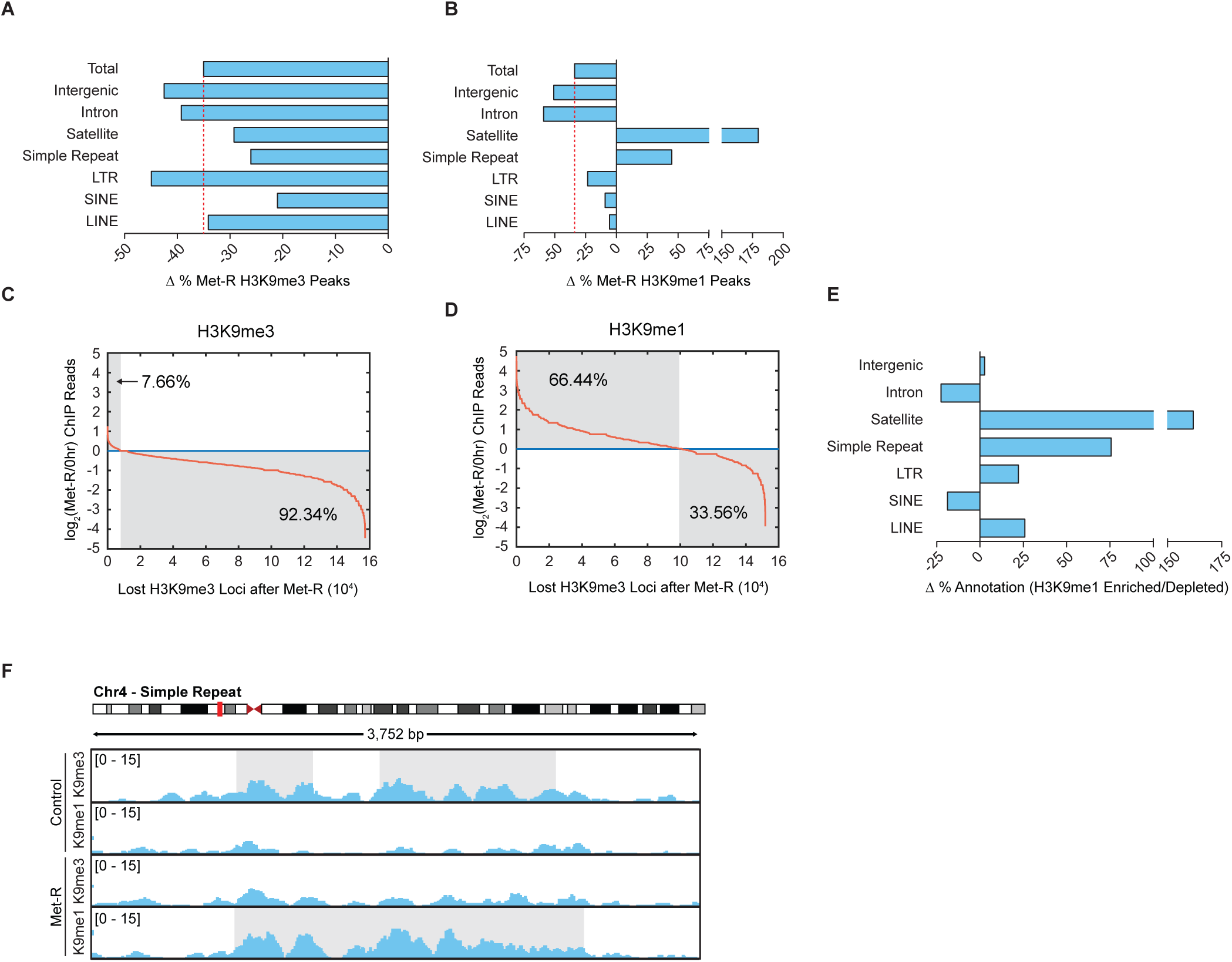
H3K9me1 is Redistributed Over Repetitive and Transposable Elements During SAM Depletion (*A-B*) Bar graphs illustrating percent changes in H3K9 PTM peak number at various annotated loci after 24 hours of Met-restriction in HCT116 cells. (*C-D*) Plots depicting changes in normalized H3K9 PTM ChIP-sequencing reads to loci of lost H3K9me3 after 24 hours of Met-restriction in HCT116 cells. (*E*) Bar graphs illustrating percent changes in loci annotation for regions of lost H3K9me3 experiencing increased relative to decreased H3K9me1 ChIP read enrichment after 24 hours of Met-restriction in HCT116 cells. (*F*) Representative ChIP-sequencing track image.

Maintained or increased levels of H3K9me1 at constitutively repressed regions during SAM depletion could result from the replacement of H3K9me1 for lost H3K9me3. To investigate this possibility, H3K9me1 reads were mapped to nearly 16e_4_ regions determined to lose H3K9me3 peak enrichment upon SAM depletion. This analysis determined H3K9me1 enrichment increases at 66.4% of these sites (Figure 4C-4D). A higher percentage of these H3K9me1 enriched sites were annotated for LINE, LTR, simple repeat, and satellite elements relative to those experiencing decreased H3K9me1 read enrichment (Figure 4E). Figure 4F provides a representative image of H3K9me3 replacement by H3K9me1 at a pericentric, repetitive loci. Together, these ChIP-sequencing analyses suggest SAM depletion stimulates the redistribution and targeting of H3K9me1 methylation patterns to repetitive and transposable genomic loci which are susceptible to higher state H3K9 demethylation.

### *De novo* H3K9 Mono-methylation Preserves Heterochromatin Stability

Replacement of H3K9me3 by H3K9me1 may function to preserve heterochromatin stability and pericentric chromatin repression during SAM depletion by providing the primer for site-specific H3K9me2/3 methylation and/or by preventing H3K9 acetylation. An MNase accessibility assay was used to evaluate global heterochromatin stability under SAM depleted conditions as MNase can more readily digest euchromatic DNA compared to heterochromatic DNA. Coupling Met-restriction with UNC0642 treatment resulted in a significantly greater loss and accumulation of di- and mono-nucleosome species, respectively, compared to Met-restriction alone (Figure 5A-5B). This indicates inhibition of *de novo* H3K9 mono-methylation during SAM depletion has a more detrimental effect on global heterochromatin stability than SAM depletion in isolation. To determine if global decreases in heterochromatin stability resulted in de-repression of transposable DNA elements, RT-qPCR was employed. Transcript abundance of *LINE 1*, *HERV-K*, and *HERV-R* were all significantly elevated after SAM depletion. Coupling SAM depletion with UNC0642 treatment resulted in the further elevation of *LINE1* and *HERV-K* transcript abundance while *HERV-R* expression was unaffected (Figure 5C-5E). Because SAM depletion-induced losses in heterochromatin stability and retrotransposon repression are exacerbated with inhibition of nuclear Ehmt1/Ehmt2 activity, these data suggest stimulation of *de novo* H3K9 mono-methylation functions, in part, to preserve heterochromatin stability.

**Figure 5.**
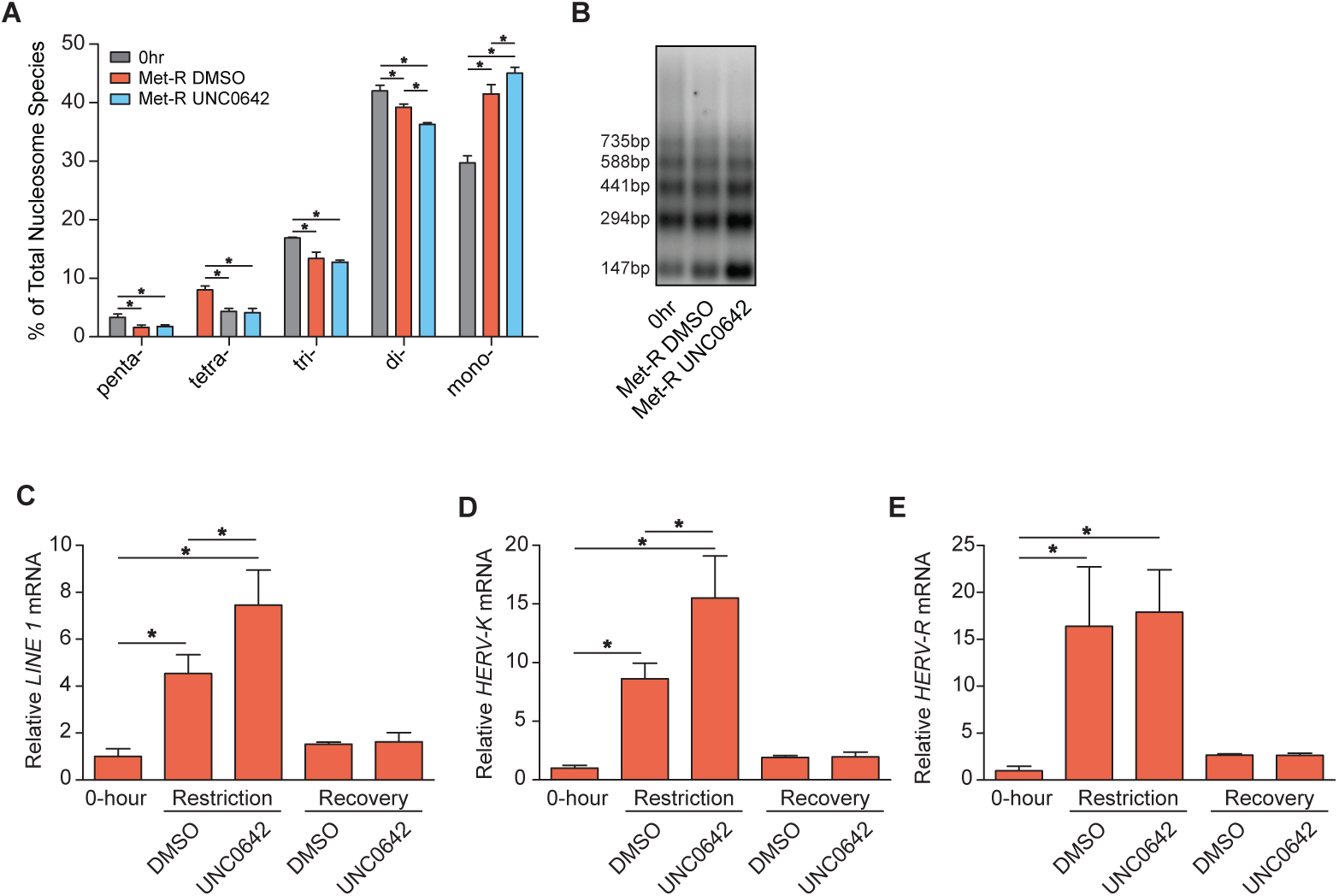
*De novo* H3K9 Mono-methylation Preserves Heterochromatin Stability (*A*) Bar graph illustrating the percent abundances of 5 distinct nucleosome species following MNase digestion of HCT116 cells. n 3, error bars represent SD, *p<0.05 (Welch’s t-Test). (*B*) Representative DNA agarose gel image of those used for the quantification presented in Figure 5A. (C*-E*) Bar graphs illustrating relative mRNA abundances of *LINE* 1, *HERV-K,* and *HERV-R* 4, error bars represent SD, *p<0.05 (Welch’s t-Test). See also Table S2.

### Adaptive H3K9 Mechanisms Support Epigenetic Persistence Upon Metabolic Recovery

Dysregulation of adaptive epigenetic mechanisms during SAM depletion may produce aberrant changes in histone PTMs that compromise long-term epigenetic persistence upon metabolic recovery. To determine if *de novo* H3K9me1 is required for epigenetic persistence to SAM depletion, HCT116 cells were allowed to recover in Met-replete media after being subjected to Met-restriction coupled with acute DMSO or UNC0642 treatment (Figure 6A). Histone proteomics analysis confirmed that inhibition of Ehmt1/Ehmt2 activity during SAM depletion results in a significant loss of H3K9me1 abundance (Figure 6B-6C). Upon 5 and 24 hours of Met-repletion, DMSO control cells largely regained their original histone PTM state (Figure 6B). However, histone PTM states of cells acutely treated with UNC0642 only during Met-restriction remained dysregulated after both 5 and 24 hours of Met-repletion (Figure 6B). Although significant global dysregulation was apparent, the H3K9 peptide proteoforms displayed the most dynamic differences (Figure 6D-6E). Interestingly, the signature H3K9 PTM response to SAM depletion described in Figure 2 was largely retained upon Met-repletion in cells acutely treated with UNC0642 (Figure 6D-6E). H3K9me3 abundance was an exception, recovering to levels consistent with regained *LINE 1* and *HERV-K/R* repression (Figure 5C-5E).

**Figure 6.**
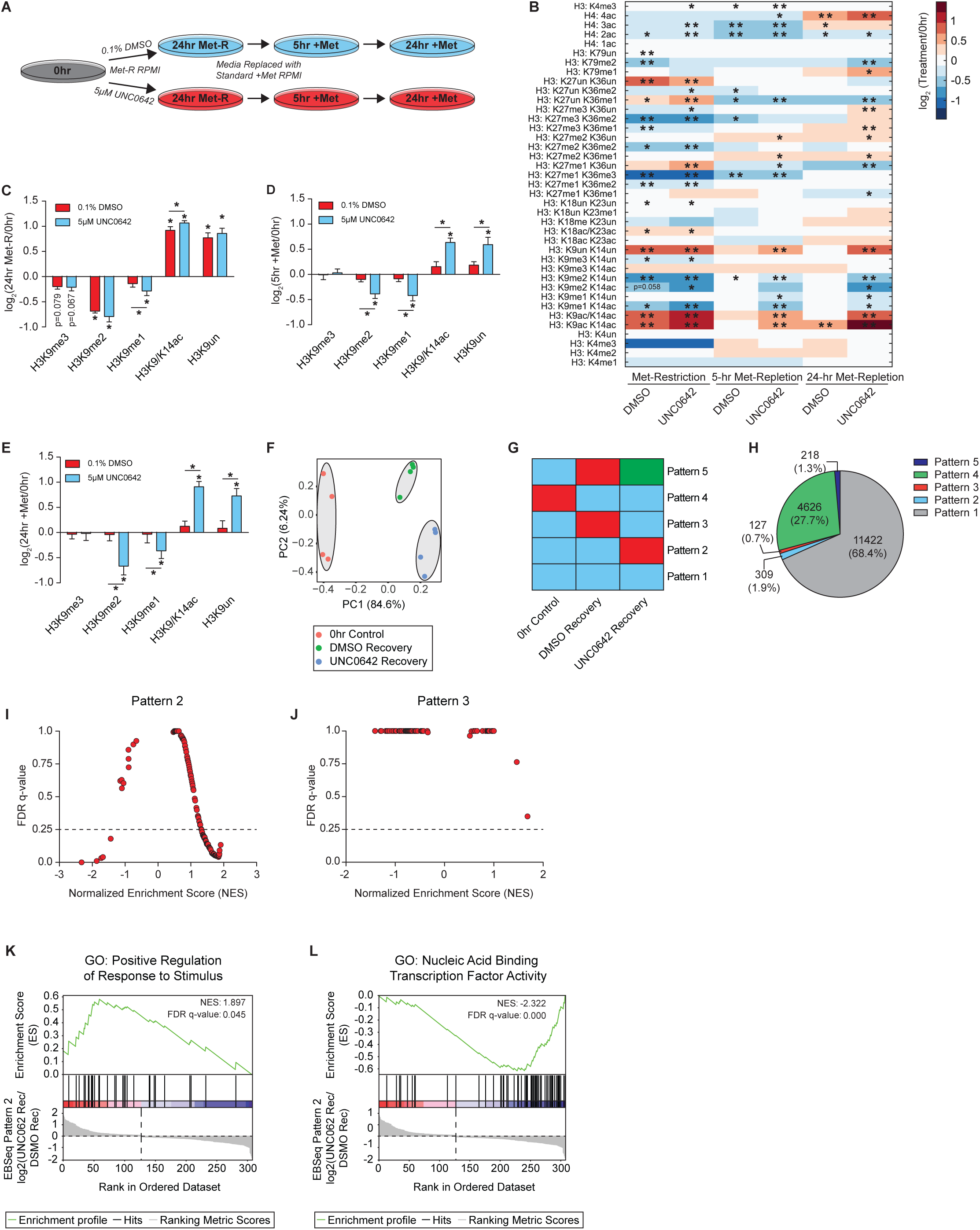
Adaptive H3K9 Mechanisms Support Epigenetic Persistence Upon Metabolic Recovery (*A*) Diagram illustrating the experimental design for replenishing Met and SAM availability after 24 hours of Met-restriction coupled with acute 5 *μ*M UNC0642 or 0.1% DMSO treatment in HCT116 cells. (*B*) Heatmap of log_2_ fold-change stoichiometric values for histone H3 and H4 peptides as measured by LC-MS/MS. n=4, *p<0.05, **p<0.01 (Welch’s t-Test). (*C-E*) Bar graphs illustrating log_2_ fold-changes for individual H3K9 PTMs in HCT116 cells as measured by LC-MS/MS. n=4, error bars represent SD, *p<0.05 (Welch’s t-Test). (*F*) PCA diagram of TPM values. (*G*) EBseq multiple condition comparison diagram. (*H*) Pie chart illustrating percent allocation of genes identified via RNA-sequencing to all EBseq patterns. (*I-J*) Scatter plots illustrating NES and FDR q-value for GSEA identified GO terms. (*K-L*) Representative GSEA running enrichment score figures. See also Table S1 and Figure S4.

Met-replete cells allowed to recover from acute UNC0642 treatment also experienced prolonged changes in H3K9 PTM abundance after inhibitor removal (Figure S4A). As Ehmt1/2 inhibition by UNC0642 was determined to be reversible (Figure SB-S4C), these results suggest disrupted persistence to small molecule treatment – in either isolation from or conjunction with SAM depletion – is a direct consequence of acute losses in Ehmt1/Ehmt2 activity.

To assess the transcriptional consequences of lost epigenetic persistence, RNA-sequencing was performed on cells which recovered in Met-replete media after Met-restriction coupled with acute DMSO or UNC0642 treatment. Principle Component Analysis (PCA) of normalized Transcripts Per Million (TPM) values produced 3 distinct clusters, suggesting altered epigenomes correlate with altered transcript profiles in these cells (Figure 6F). A multiple condition comparison analysis was used to determine which transcripts distinguished the clusters from one another (Figure 6G). While 68.4% of identified genes were ubiquitously expressed across samples (Pattern 1), 27.7% were uniquely expressed in control cells relative to both recovery groups (Pattern 4) (Figure 6H). This result supports the PCA findings as the pre-restriction control group is most distinct from both recovery groups. Unique gene expression profiles for UNC0642 (Pattern 2) and DMSO (Pattern 3) recovery cells contained 1.9% and 0.7%, respectively, of all identified genes. Therefore, these subsets of genes are responsible for the unique clustering of both Met-restriction recovery groups (Figure 6H). Gene Set Enrichment Analysis (GSEA) was performed on both Pattern 2 and 3 gene lists to determine functional differences between each gene set. GSEA of Pattern 2 genes identified 71 significantly enriched gene ontology (GO) terms possessing an FDR q-value less than 0.25, while an identical analysis of Pattern 3 genes did not identify any significantly enriched GO terms (Figure 6I-6J). Of the significantly enriched GO terms in Pattern 2, 66 possessed a positive Normalized Enrichment Score (NES) when comparing UNC0642 to DMSO control recovery cells (Figure S5A-S5B). Figure 6K is a representative GSEA running enrichment score figure of a GO term possessing a positive NES. Only 5 of the significantly enriched GO terms in Pattern 2 possessed a negative NES when comparing UNC0642 to DMSO control recovery cells. Notably, a majority of the genes assigned to these GO terms were zinc finger binding proteins (Figure S5C-S5D). Figure 6L is a representative GSEA running enrichment score figure of a GO term possessing a negative NES.

These data suggest adaptive regulation of H3K9 PTMs in response to SAM depletion is required for long-term epigenetic persistence upon metabolic recovery. Furthermore, inhibited epigenetic persistence results in significant transcriptional consequences that impact a diverse set of cellular functions.

### Adaptive and Persistent Epigenetic Responses to SAM Depletion are Robust *in vivo*, Independent of Age

To determine if adaptive and persistent responses to SAM depletion exist *in vivo* and are age-independent, 6- and 22-month C57BL/6J mice were subjected to 3 weeks of Met-restriction followed by 5 weeks of Met-repletion (Figure 7A). Several previous reports have associated decreased H3K9me2/3 abundance with aging in numerous organisms (Wood et al., 2010; Larson et al., 2012; Ni et al., 2012; Tvardovskey et al., 2017; Scaffidi and Misteli, 2006). Age-dependent dysregulation of heterochromatin has been associated with de-repression of repetitive DNA and transposable elements, as well as DNA damage, factors which may contribute to disease progression (Wood et al., 2016; De Cecco et al., 2013; De Cecco et al., 2019; Oberdoerffer and Sinclair, 2007; Booth and Brunet, 2016). Therefore, conservation of both adaptive and persistent responses to SAM depletion across the lifespan would further support the functional importance of both mechanisms as an “older” chromatin environment is thought to be inherently dysregulated.

**Figure 7.**
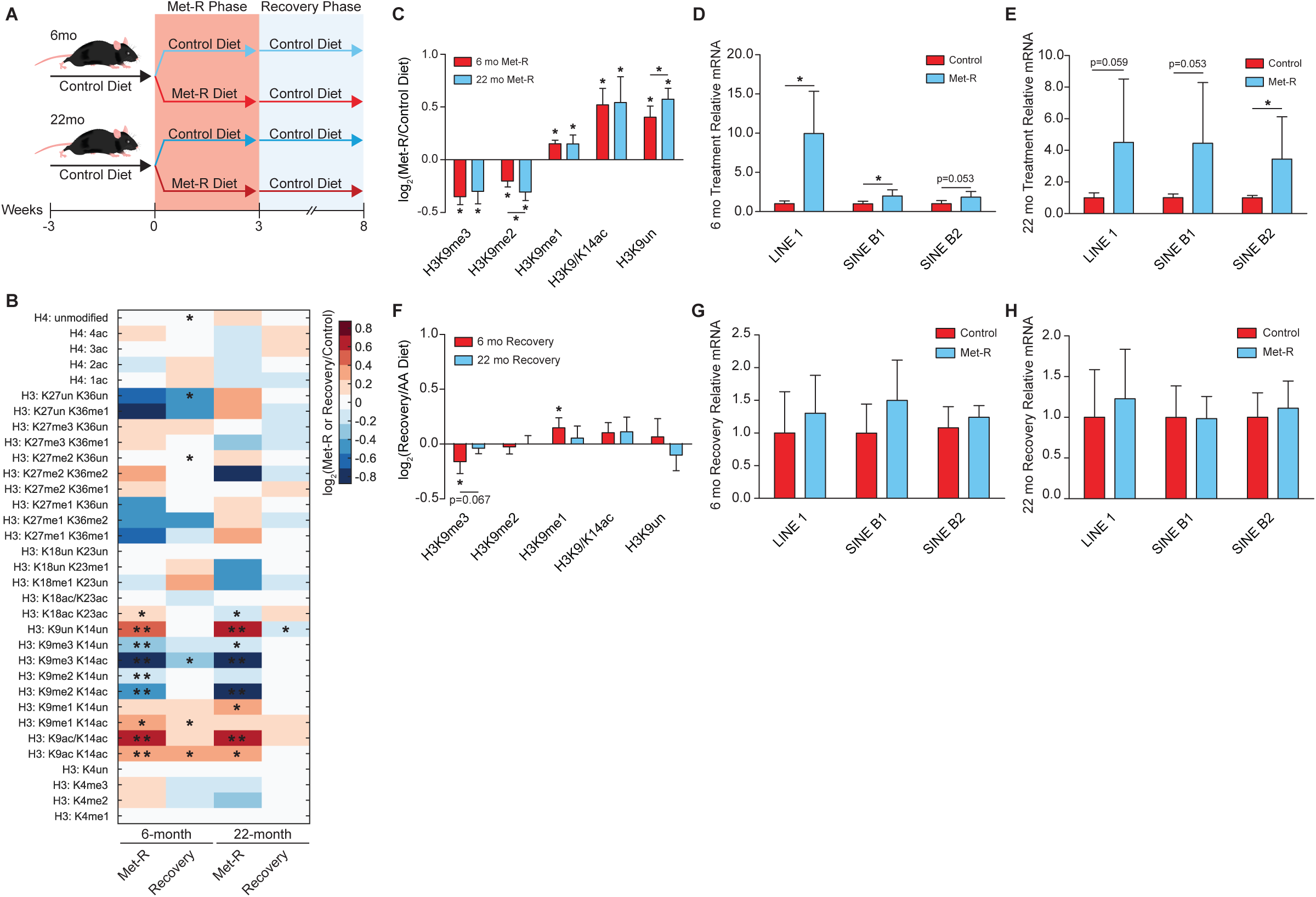
Adaptive and Persistent Epigenetic Responses to SAM Depletion are Robust *in vivo*, Independent of Age (*A*) Diagram depicting the experimental design for *in vivo* Met-restriction of 6- and 22-month C57BL/6J mice followed by dietary Met-reintroduction. (*B*) Heatmap of log_2_ fold-change stoichiometric values for histone H3 and H4 peptides in C57BL/6J liver relative to age-matched controls as measured by LC-MS/MS. n 5, *p<0.05, **p<0.01 (Welch’s t-Test). (*C*) Bar graph illustrating log_2_ fold-changes for individual H3K9 PTMs in liver from Met-restricted C57BL/6J 5, error bars represent SD, *p<0.05 (Welch’s t-Test). (*D-E*) Bar graphs illustrating relative mRNA abundances of *LINE 1*, *SINE B1*, and *SINE B2* in C57BL/6J liver as measured by RT-qPCR. n 5, error bars represent SD, *p<0.05 (Student’s t-Test). (*F*) Bar graph illustrating log_2_ fold-changes for individual H3K9 PTMs in liver from C57BL/6J relative to age-matched controls as measured by 5, error bars represent SD, *p<0.05, (Welch’s t-Test). (*G-H*) Bar graphs illustrating relative mRNA abundances of *LINE 1*, *SINE B1*, and *SINE B2* in C57BL/6J liver as 5, error bars represent SD, *p<0.05 (Student’s t-Test). See also Table S1, Table S2, and Figure S5.

Remarkably, histone proteomics analysis of liver tissue revealed H3K9 was the only residue on which PTMs responded similarly to SAM depletion in both 6- and 22-month mice relative to age-matched controls (Figure 7B). This PTM response at H3K9 phenocopied those identified in the previously described *in vitro* and whole-organism SAM depletion studies, further highlighting the robust regulation of this residue (Figure 7C). Significant age-dependent responses to SAM depletion were identified on all remaining histone residues (Figure 7B and S6A-S6E). Furthermore, significantly elevated *LINE 1*, *SINE B1*, and *SINE B2* transcript abundance in both 6- and 22-month mice suggest the chromatin instability phenotype characterized in Figure 5 also occurs during metabolic depletion of SAM *in vivo* (Figure 7D-7E). After 5 weeks of dietary Met-reintroduction, both young and old mice displayed the ability to re-establish an epigenome similar to that possessed by age-matched controls (Figure 7B). Interestingly, 22-month mice appeared to more rapidly adopt the control histone PTM state, while the 6-month mice still exhibited residual losses in H3K9me3 and increases in H3K9me1 after 5 weeks of recovery (Figure 7F). Consistent with the general recovery of chromatin states, both age groups largely reestablished repression of *LINE 1*, *SINE B1*, and *SINE B2* expression (Figure 7G-7H).

Metabolic changes in young and old mice tracked with H3K9 PTMs. Both 6- and 22-month mice in the Met-restriction group experienced significant losses in total body weight relative to age matched controls (Figure S6F). Decreases in overall body weight were comprised of significant losses in both lean and fat mass (Figure S6F). Altered body-weight and compositions were accompanied by significantly improved glucose tolerance in both young and old mice (Figure S6G). Upon Met-repletion, mice reacquired their initial metabolic state in an age-independent manner. All animals experienced significant increases in total body weight, comprised of elevated lean and fat mass, as well normalized glucose tolerance relative to age-matched controls (Figure S6H-S6I).

Together, strong association of the H3K9 chromatin response to SAM depletion with metabolic reprogramming in young and old mice suggest H3K9 regulatory mechanisms are not only robust across lifespan but may be required to support these metabolic phenotypes. Furthermore, these data show the ability of epigenetic states to persist upon recovery from a metabolic challenge is an inherent property of both isolated cells and complex mammals.

## Discussion

Here, we provide a comprehensive investigation into the adaptive regulation of chromatin during a metabolic stress. A highly conserved, robust histone methylation response to SAM depletion was discovered, characterized by global losses in histone H3 di- and tri-methylation while H3K9me1 is actively maintained to safeguard against exacerbated heterochromatin instability and support epigenetic persistence upon metabolic recovery. These mechanisms involve the enrichment of H3K9me1 in susceptible regions of heterochromatin through *de novo* methylation of both chromatin-bound and cytoplasmic H3. To our knowledge, this is the first identified mechanism in which scarce, diet-derived nutrients are funneled toward the preservation of specific chromatin PTM states.

Zee et al., 2010 have previously determined H3K9me1 has the shortest half-life of 17 measured PTMs, highlighting the importance of *de novo* methylation for supporting global H3K9me1 abundance in a SAM depleted environment. Here, *de novo* methylation was found to be critical for more than simply sustaining global H3K9me1 levels under these conditions. ChIP-sequencing analyses determined H3K9me1 patterns experience a significant redistribution across heterochromatin that results in H3K9me1 enrichment at repetitive and transposable DNA elements, particularly in regions where H3K9me3 is lost. Targeting of H3K9me1 to these regions is likely critical for preserving heterochromatin as prevention of *de novo* H3K9 mono-methylation resulted in global losses in heterochromatin stability and site-specific retrotransposon derepression. We hypothesize H3K9me1 facilitates repression of retrotransposons during SAM depletion through two simultaneous mechanisms: 1.) acting as the primed substrate for HMTs which deposit di- and tri-methyl PTMs onto H3K9 (Peters et al., 2001; Loyola et al., 2006; Loyola et al., 2009; Rea et al., 2000; Bannister et al., 2001; Lachner et al., 2000) and 2.) preventing the acetylation of newly demethylated H3K9 residues. Increased H3K9 acetylation through impaired H3K9 deacetylation pathways is associated with global genomic instability as well as derepression of LINE1, providing support for the second mechanism (Mostoslavsky et al., 2006; Van Meter et al., 2014; Simon et al., 2019). Ensuring sufficient amounts of H3K9me1 are available to facilitate heterochromatin stability is not limited to nuclear derived H3K9me1. Pinheiro et al., 2012 have previously shown inhibition of cytoplasmic H3K9 mono-methylation through shRNA knockdown of *PRDM3* and *PRDM16* stimulates pericentric heterochromatin loss and disruption of the nuclear lamina. Our study demonstrates cytoplasmic derived H3K9me1 comprises a significant portion (18%) of the total chromatin bound pool after 24 hours of Met-restriction *in vitro*. Adaptation of H3K9 mono-methylation to SAM depletion is also critical for long-term epigenetic persistence to this metabolic stress. Prevention of *de novo* H3K9 mono-methylation during SAM depletion resulted in sustained dysregulation of global histone PTMs upon metabolic recovery, significantly influencing gene expression patterns. This requirement for an active, adaptive histone PTM response to support epigenetic persistence upon metabolic recovery is the first identified mechanism of its kind.

Replication of *in vitro* results *in vivo*, independent of age, strongly supports the discovery of a universal adaptive and persistent response to SAM depletion. Here, we demonstrate that the adaptive mechanisms required to support H3K9me1-mediated heterochromatin stability are inducible in both 6- and 22-month mice after dietary Met-restriction. Furthermore, H3K9me1 maintenance under SAM depleted conditions uniquely correlated with beneficial metabolic reprogramming (i.e. reduced fat mass and improved glucose tolerance) in both young and old mice. This suggests the robust regulation of H3K9 PTMs in response to SAM depletion may be required to support the metabolic reprogramming stimulated by dietary Met-restriction. Global epigenetic persistence to metabolic SAM depletion was observed in both 6- and 22-month mice. Validation of this phenomenon *in vivo*, independent of age, suggests similar mechanisms critical for maintaining proper regulation of the epigenome during life-experienced fluctuations in the availability of other metabolic cofactors may exist as well. Such fluctuations could be stimulated by periods of prolonged fasts, chronic intake of foods lacking essential cofactors (i.e. vegan and vegetarian dietary patterns), and circadian regulated changes in metabolite availability (Farmer, 2014; Schmidt et al., 2016; Krishnaiah et al., 2017).

Notably, we did not observe an age-associated decrease in H3K9 methylation between 6- and 22-month control diet mice (Table S1). Similar regulation of H3K9 PTMs and mirrored metabolic responses to SAM depletion across age groups suggest our 22-month mice, the equivalent to a 60-70 year old human, have not yet experienced a decline in health-span (Flurkey et al., 2007). It will be interesting to determine if the adaptive and persistent epigenetic mechanisms revealed here are functional in very old animals (i.e. >30 months) where H3K9 methylation is thought to be inherently dysregulated. Disruption of these mechanisms later in an organism’s lifespan could support age-associated derepression of transposable elements (Wood et al., 2016; De Cecco et al., 2013; Oberdoerffer and Sinclair, 2007; Booth and Brunet, 2016), contributing to negative phenotypes that accelerate the aging process such as inflammation (De Cecco et al., 2019; Simon et al., 2019).

Future studies will be needed to determine how H3K9 mono-methyltransferases are able to maintain their catalytic activity in SAM depleted environments. Favorable steady-state kinetic constants for H3K9me1 relative to H3K9me2/3 HMTs is one possibility. However, it is difficult to address this hypothesis by comparing existing literature on H3K9 HMT enzymology, due to disparate substrates and assays. Direct shuttling of SAM to H3K9me1 HMTs by Mat2A could also support H3K9me1 HMT catalytic activity. As SAM is energetically expensive to produce, due to the 1:1 requirement of Met and ATP molecules for its synthesis, in addition to being highly unstable (Wu et al., 1983), it would be advantageous for the cell to possess mechanisms which permit preferential shuttling of SAM during methyl-metabolite depletion. Directed SAM production and utilization have been proposed in other metabolic contexts (Katoh et al., 2011; Kera et al., 2013; Li et al., 2015).

## Acknowledgements

This research was supported in part by grants from the NIH (S.A.H. –T32 DK007665; K.A.K. T32 DK007665; W.H.L. T32 DK007665; J.M.D. – R37 GM059785; D.W.L. – AG041765, AG050135, AG051974, AG056771, AG062328, a New Investigator Program Award (D.W.L.) and a Collaborative Health Sciences Program Award (V.L.C.) from the Wisconsin Partnership Program, the V. Foundation for Cancer Research (V.L.C.), and a Glenn Foundation Award for Research in the Biological Mechanisms of Aging (D.W.L.), as well as startup funds from the UW-Madison School of Medicine and Public Health and the UW-Madison Department of Medicine (V.L.C. and D.W.L.). This research was conducted while D.W.L. was an AFAR Research Grant recipient from the American Federation for Aging Research. D.Y. is supported in part by a fellowship from the American Heart Association (17PRE33410983). The Lamming laboratory is supported in part by the U.S. Department of Veterans Affairs (I01-BX004031), and this work was supported using facilities and resources from the William S. Middleton Memorial Veterans Hospital. This work does not represent the views of the Department of Veterans Affairs or the United States Government. We would like to thank Alexis J. Lawton for the bioinformatics analysis presented in Figure S3I. We would also like to thank the University of Wisconsin-Madison Biotechnology Center and the University of Wisconsin-Madison Center for High Throughput Computing for their help in generating and analyzing the ChIP-sequencing data presented in Figure 5, respectively.

## Author Contributions

Conceptualization, S.A.H., D.W.L, V.L.C., and J.M.D.; Methodology, S.A.H., C.Y., C.K.W., L.N.C., W.H.L., K.I., B.P.T., D.W.L., and J.M.D.; Validation, S.A.H., D.Y., C.Y., L.N.C., K.A.K., K.I., B.P.T., D.W.L., and J.M.D.; Formal Analysis, S.A.H., D.Y., C.K.W., K.A.K., D.W.L., and J.M.D; Investigation, S.A.H., D.Y., C.Y., L.N.C., K.A.K., J.L.T., S.E.Y., and B.R.M.; Resources, K.I., B.P.T., V.L.C., D.W.L., and J.M.D.; Writing – Original Draft, S.A.H. and J.M.D.; Writing – Review & Editing, S.A.H., D.Y., C.Y., C.K.W., W.H.L., K.I., R.S., B.P.T., V.L.C., D.W.L., and J.M.D.; Visualization, S.A.H., D.Y., and D.W.L.; Supervision and Project Administration, K.I., R.S., B.P.T., D.W.L., and J.M.D.; Funding Acquisition, S.A.H., K.I., R.S., B.P.T., V.L.C., D.W.L., and J.M.D.

## Supplemental Figure Titles and Legends

**Figure S1, Related to Figure 1.**
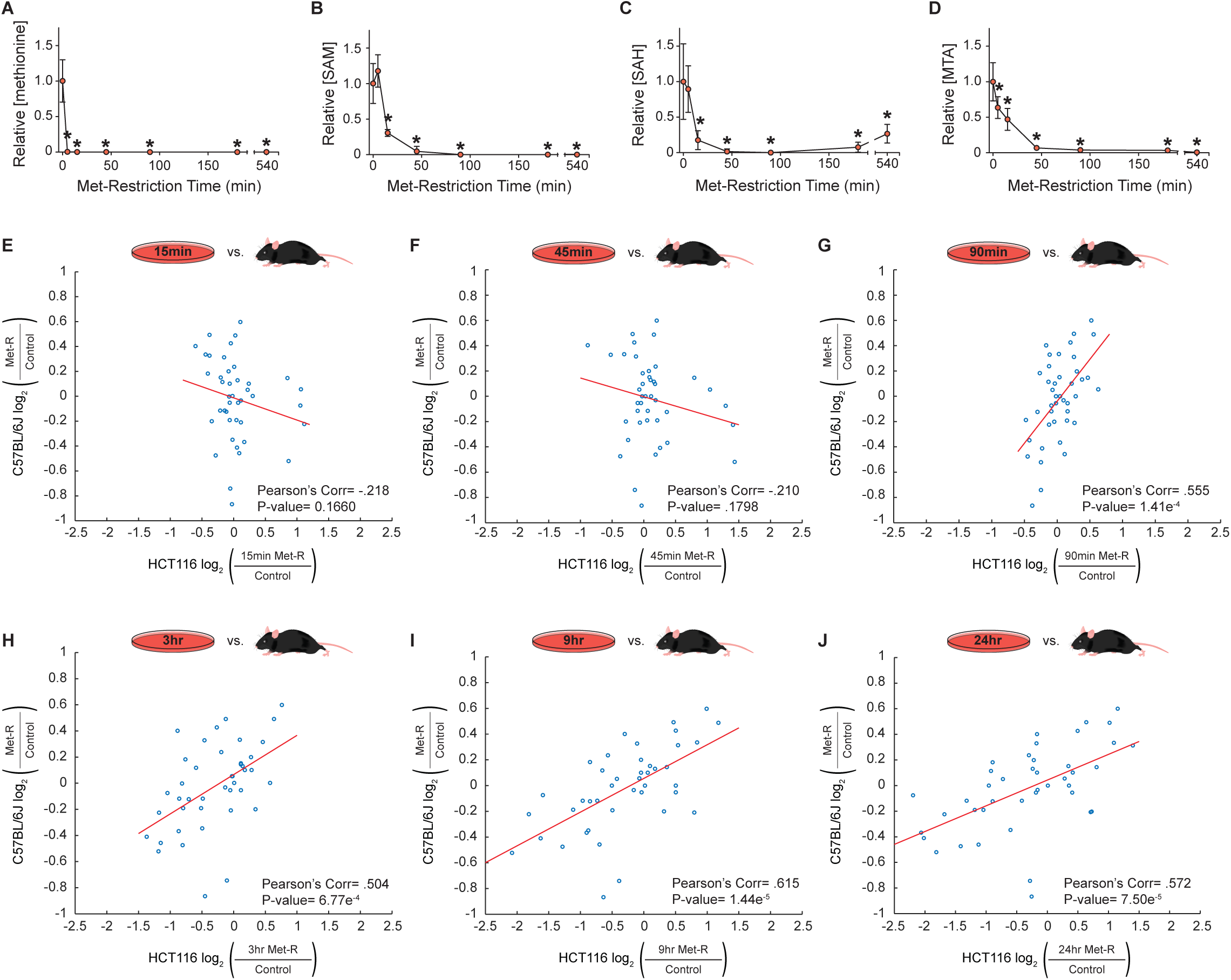
(*A-D*) Scatter plots of relative abundance values for key Met-cycle metabolites in HCT116 cells as measured by LC-MS. n=5, error bars represent SD, *p<0.05 (Welch’s t-Test). (*E-J*) Correlation plots of LC-MS/MS generated log_2_ fold-change stoichiometric values for individual histone H3 peptide proteoforms. n≥3. See also Table S1.

**Figure S2, Related to Figure 2.**
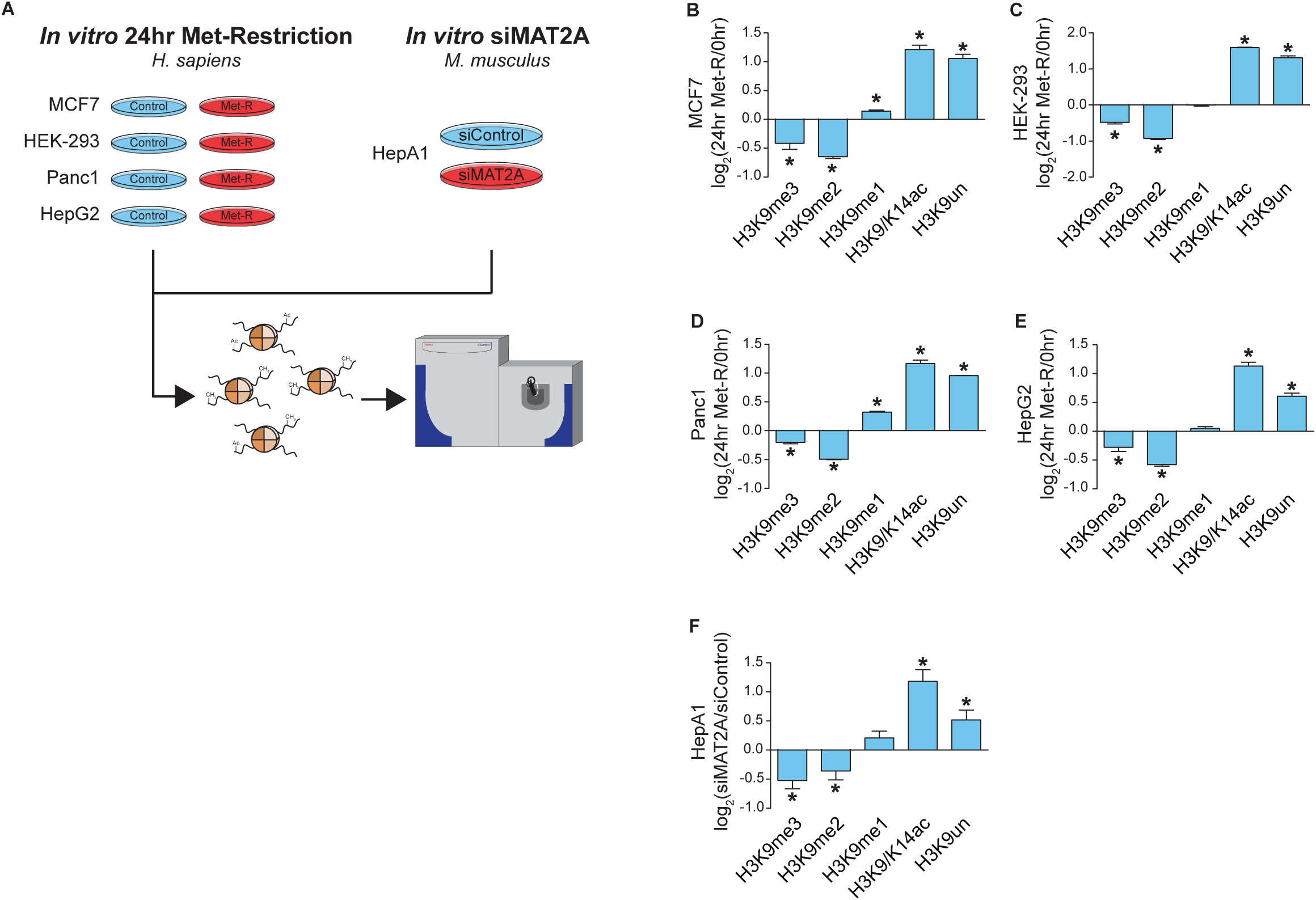
(A) Diagram illustrating the experimental design for determining the conservation of H3K9 regulation in response to SAM depletion across various biological systems. (*B-F*) Bar graphs illustrating log_2_ fold-changes for individual H3K9 in response to *in vitro* SAM depletion. n=3, error bars represent SD, *p<0.05 (Welch’s t-Test). See also Table S1.

**Figure S3, Related to Figure 3.**
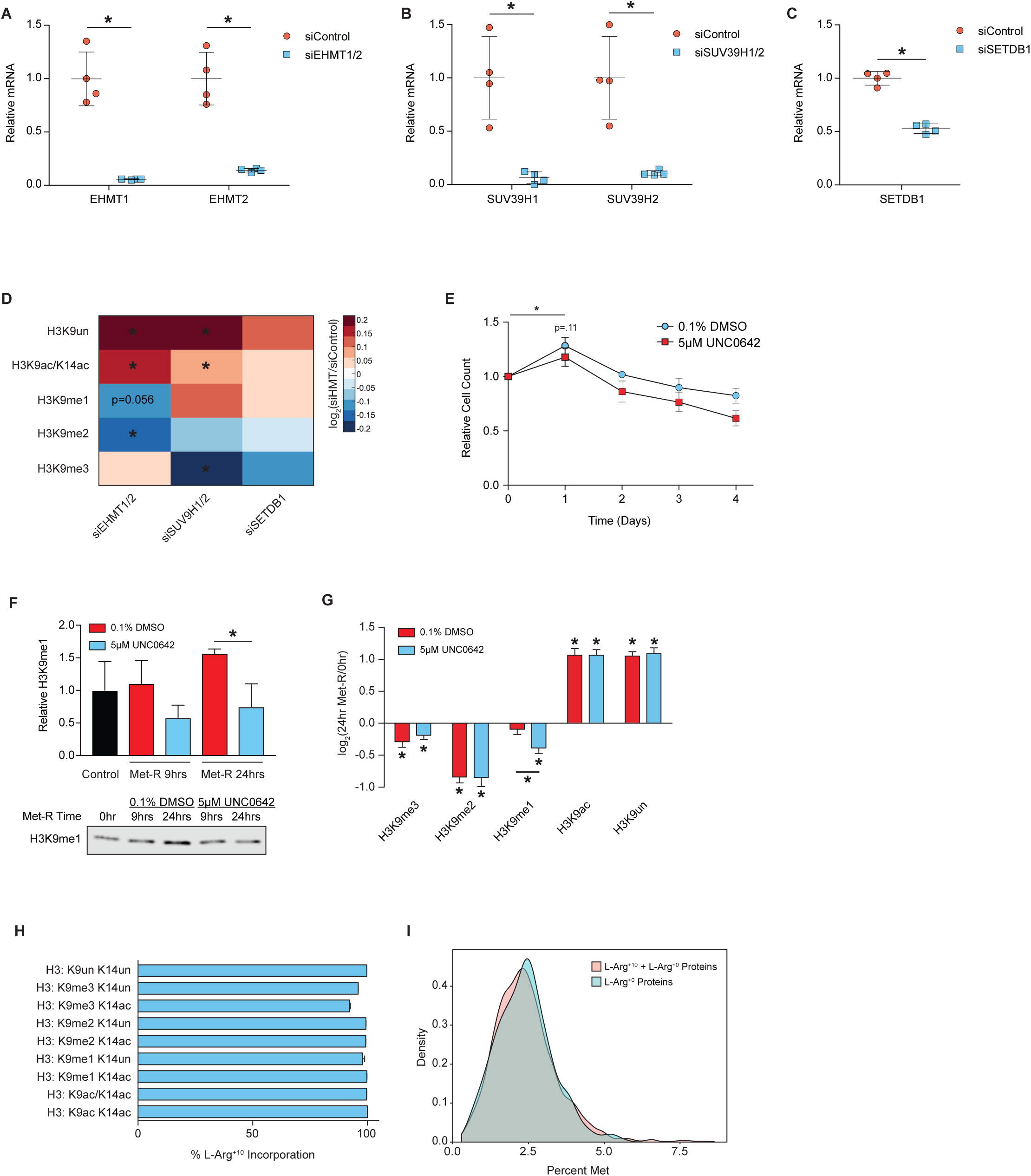
(*A-C*) Scatter plots of relative HMT mRNA abundance in HCT116 cells as measured by RT-qPCR. n, error bars represent SD, *p<0.05 (Welch’s t-Test). (*D*) Heatmap of log_2_ fold-change stoichiometric values for unique histone H3K9 peptide proteoforms in HCT116 cells. n=4, *p<0.05 (Welch’s t-Test). (*E*) Scatter plot of relative cell counts for attached HCT116 cells. n=4, error bars represent SD, *p<0.05 (Welch’s t-Test). (F) Bar graph of relative H3K9me1 abundance in HCT116 cells as measured by western blot with accompanied representative blot image. See Document S1 for REVERT Total Protein Stain image used for normalization. Error 3, *p<0.05 (Welch’s t-Test). (*G*) Bar graphs illustrating log_2_ fold-changes for individual H3K9 PTMs. n 3, *p<0.05 (Welch’s t-Test). (*H*) Bar graph showing average incorporation of L-Arg_+10_ in H3K9 peptide proteoforms. n=4, error bars represent SD. (*I*) Density plot illustrating the percentage of Met residue frequency in the primary amino acid sequence of all identified cytoplasmic proteins as well as those determined to be preferentially translated. See also Table S1, Table S2, and Table S3.

**Figure S4, Related to Figure 6.**
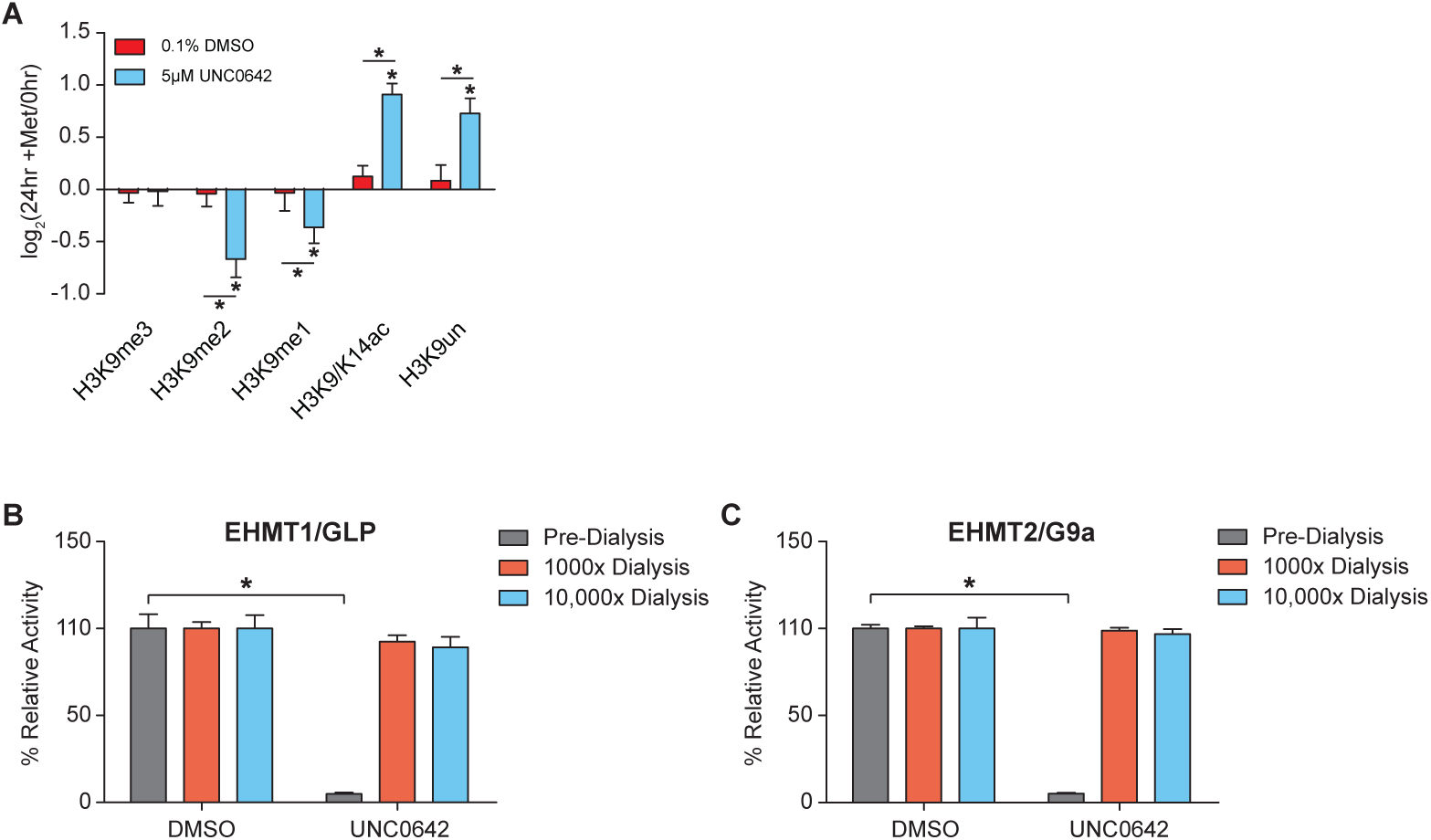
(*A*) Bar graphs illustrating log_2_ fold-changes for individual H3K9 PTMs 24 hours following UNC0642 removal. n=4, error bars represent SD, *p<0.05 (Welch’s t-Test). *(B-C)* Bar graphs illustrating relative recombinant enzyme activity in the presence of saturating amounts UNC0642 and following 1000x or 10,000x overnight dialysis. n=4, error bars represent SD, *p<0.05 (Welch’s t-Test).

**Figure S5, Related to Figure 6.**
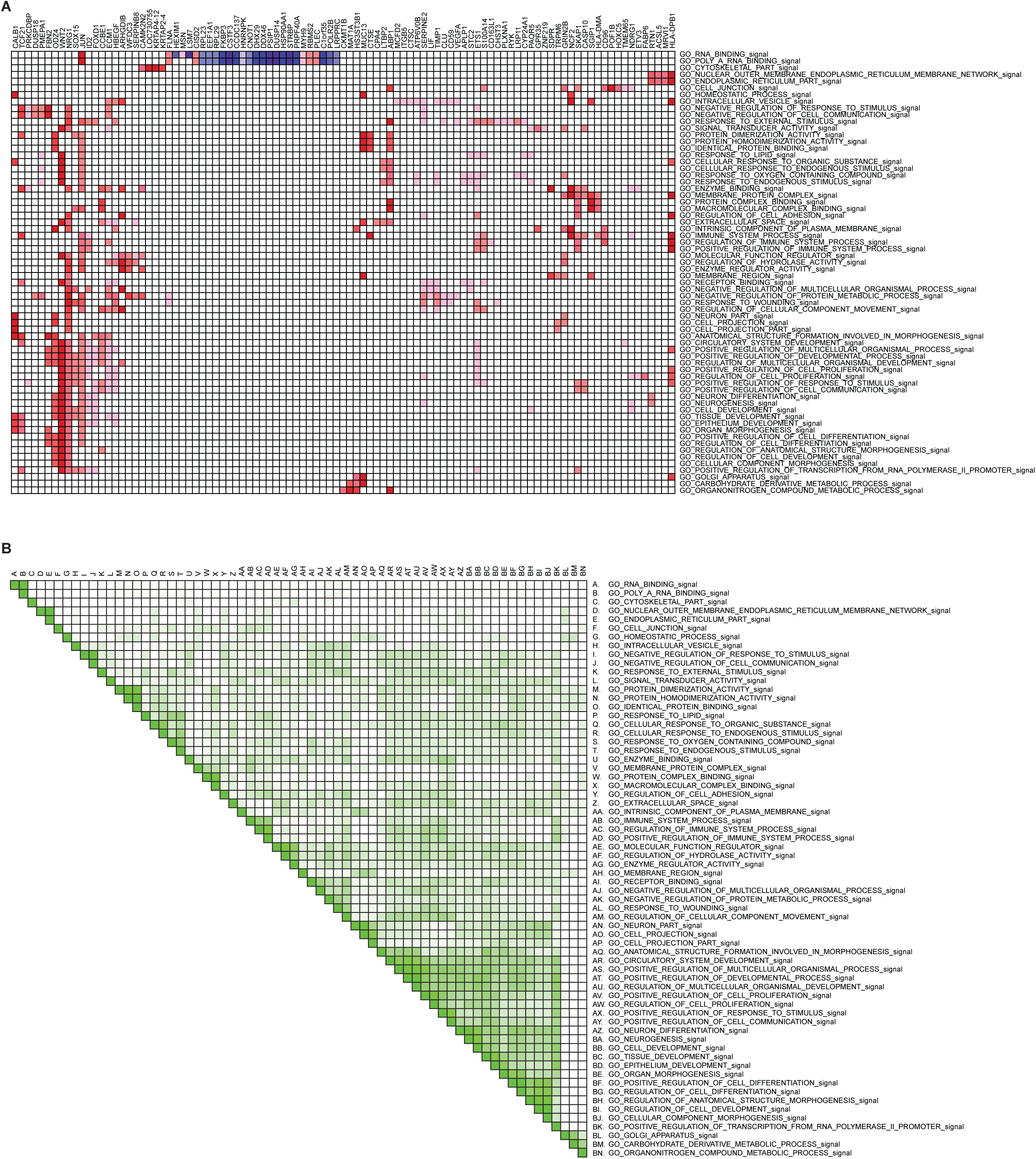
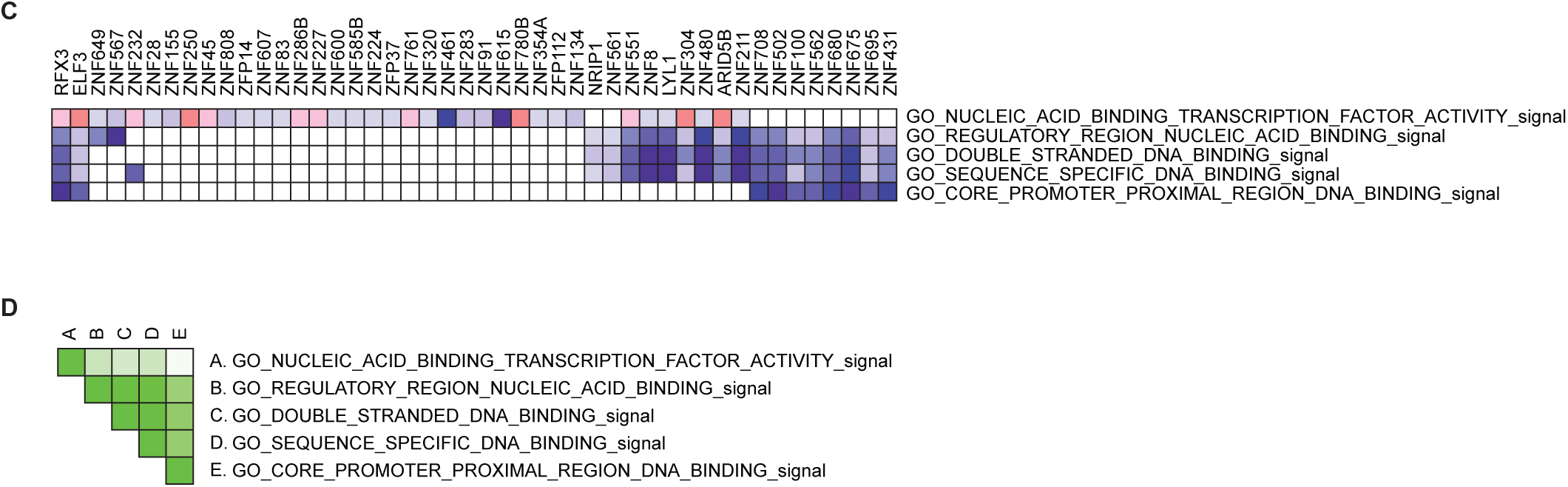
(*A*) GSEA heatmap of relative gene expression across positive NES GO terms for Pattern 2. (B) GSEA set-to-set heatmap illustrating gene enrichment overlap across positive NES GO terms for Pattern 2. (*C*) GSEA heatmap of relative gene expression across negative NES GO terms for Pattern 2. (D) GSEA set-to-set heatmap illustrating gene enrichment overlap across negative NES GO terms for Pattern 2.

**Figure S6, Related to Figure 7.**
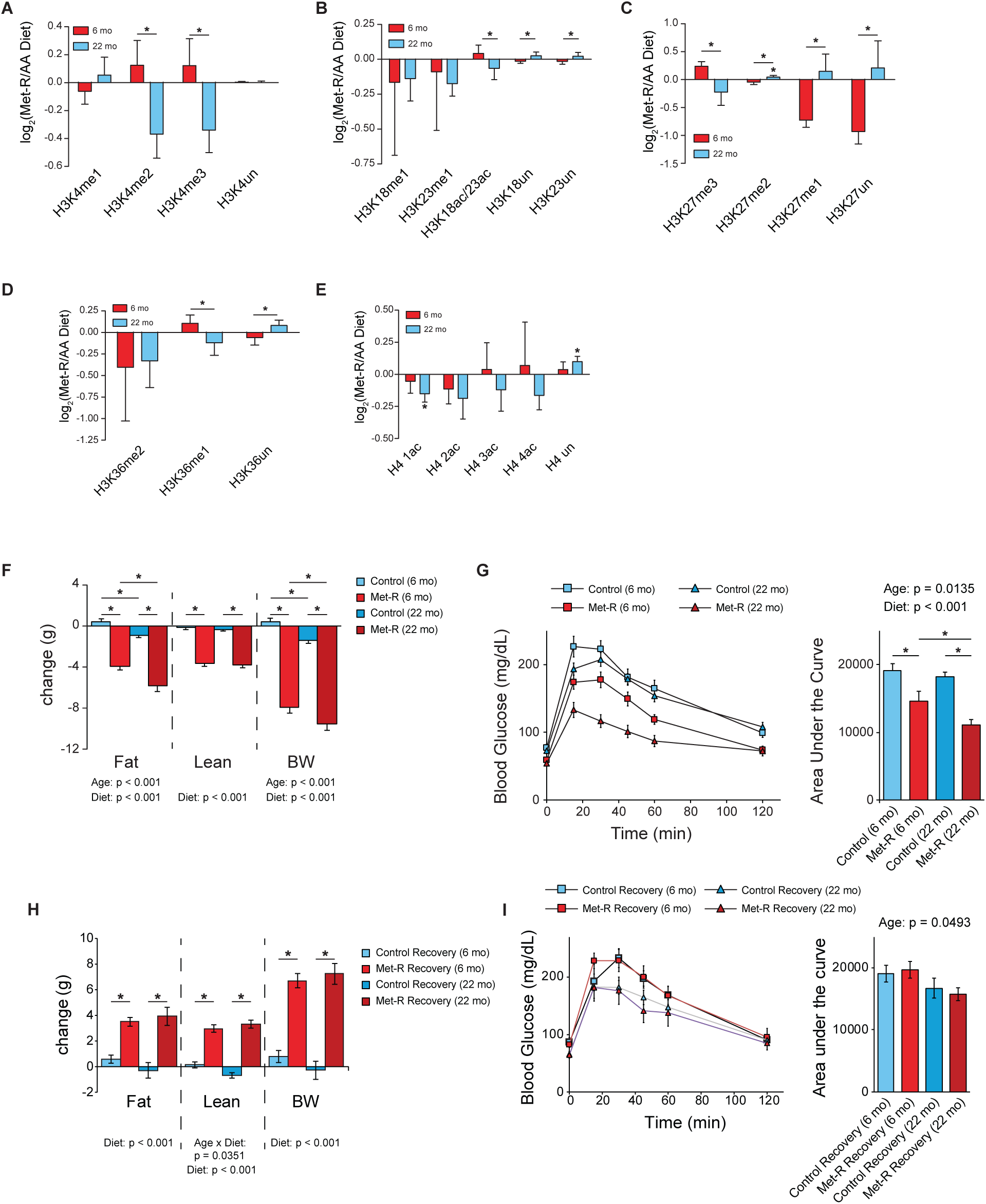
(*A-E*) Bar graphs illustrating log_2_ fold-changes for individual H3K4, H3K18/K23, H3K27, H3K36, and H4 PTMs in liver from Met-restricted C57BL/6J mice relative to age-matched controls as measured by LC-MS/MS. n≥5, error bars represent SD, *p<0.05 (Welch’s t-Test). (F) Bar graph illustrating changes in fat, lean mass, and total body mass (BW) after 3 weeks of control or Met-restricted diet feeding. n=14, error bars represent SEM, *p<0.05 (two-way ANOVA). (*G*) Glucose tolerance test in 6- and 22-month C57BL/6J mice after 2 weeks of control or Met-restricted diet feeding. n=14, error bars represent SEM, *p<0.05 (two-way ANOVA). (*H*) Bar graph illustrating changes in fat, lean mass, and total body mass (BW) after 5 weeks of control diet feeding following either 3 weeks of control or Met-restricted diet feeding. n represent SEM, *p<0.05 (two-way ANOVA). (*I*) Glucose tolerance test in 6- and 22-month C57BL/6J mice after 4 weeks of control diet feeding following either 3 weeks of control or Met-restricted diet feeding. n≥7, error bars represent SEM, *p<0.05 (two-way ANOVA). See also Table S1 and Table S2.

## Star Methods Text

### Contact for Reagent and Resource Sharing

Further information and requests for resources and reagents should be directed to and will be fulfilled by the Lead Contact, John M. Denu.

### Experimental Model and Subject Details

#### Animals and Diets

All experiments involving animals were approved by the Institutional Animal Care and Use Committee of the William S. Middleton Memorial Veterans Hospital, Madison WI. To test the effect of Met deprivation in the context of young mice, 9-week-old male C57BL/6J mice were purchased from The Jackson Laboratory (Bar Harbor, ME, USA) and placed on either an amino acid defined Control diet (TD.01084, Envigo, Madison, WI, USA) or a Met-deficient diet (TD.140119, Envigo, Madison, WI, USA) their respective diets at 14 weeks of age; full diet compositions for these diets have been previously described (PMID: 29401631). After 5 weeks on the diet, mice were sacrificed following an overnight fast and liver was flash frozen for analysis as described.

To compare the effect of Met restriction on young and old mice, C57BL/6J. Nia male mice were obtained from the National Institute on Aging Aged Rodent Colony, and housed in our animal facility for 3-4 months until reaching 5 and 21 months of age. All animals were then placed on the Control diet (TD.01084) for 4 weeks, until the mice were 6 and 22 months of age. Mice were then randomized to either remain on the Control diet, or were placed on the Met-deficient diet (TD.140119) for three weeks. Liver was collected and flash-frozen in liquid nitrogen harvested from mice euthanized after an approximately 16-hour fast.

#### Cell Lines

HCT116, HEK-293, HEK-293T, HMCF7, Panc1, HepG2, and HepA1 cell lines were cultured at 37°C with 5.0% CO_2_.

### Method Details

#### Metabolite Extraction

For HCT116 cells, ∼1.5×10^6 cells were quickly rinsed 3 times with 1.5 ml ice-cold PBS pH 7.4 before the addition of 1.5 ml of −80°C 80:20 MeOH:H_2_O extraction solvent. Cells were incubated with extraction solvent at −80°C for 15 minutes. Next, cells were scraped off their dish and transferred into a 2 ml microcentrifuge Eppendorf tube which was followed by a 5-minute, maximum speed centrifugation at 4°C. The supernatant was transferred to a new 2 ml microcentrifuge Eppendorf tube. Next, 0.5 ml extraction solvent was added to the remaining pellet, vortexed, and again centrifugated at maximum speed for 5 minutes at 4°C. The 2 supernatants from each extraction were pooled and dried completely using a Thermo Fisher Savant ISS110 SpeedVac. Dried metabolite extracts were resuspended in 80 *μ*l of 85% acetonitrile following microcentrifugation for 5 minutes at maximum speed at 4°C to pellet any remaining insoluble debris. The supernatant was then transferred to a glass vial for LC-MS analysis.

For *M. musculus* liver tissue, an average tissue weight of 75 mg was powdered in liquid nitrogen using a mortar and pestle. Powdered tissue was transferred to an individual 1.5 ml microcentrifuge Eppendorf tube and incubated with 1 ml −80°C 80:20 MeOH:H_2_O extraction solvent on dry ice for 5 minutes post-vortexing. Tissue homogenate was centrifugated at maximum speed for 5 minutes at 4°C. Supernatant was transferred to a 15 ml tube after which the remaining pellet was resuspended in 0.8 ml −20°C 40:40:20 ACN:MeOH:H_2_O extraction solvent and incubated on ice for 5 minutes. Tissue homogenate was again centrifugated at maximum speed for 5 minutes at 4°C after which the supernatant was pooled with the previously isolated metabolite fraction. The 40:40:20 ACN:MeOH:H_2_O extraction was then repeated as previously described. Next, 550 *μ*l of pooled metabolite extract for each sample was transferred to a 1.5 ml microcentrifuge Eppendorf tube and completely dried using a Thermo Fisher Savant ISS110 SpeedVac. Dried metabolite extracts were resuspended in 450 *μ*l of 85% ACN following microcentrifugation for 5 minutes at maximum speed at 4°C to pellet any remaining insoluble debris. Supernatant was then transferred to a glass vial for LC-MS analysis.

#### LC-MS Metabolite Analysis

Each prepared metabolite sample was injected onto a Thermo Fisher Scientific Vanquish UHPLC with a Waters XBridge BEH Amide column (100 m x 2.1 mm, 3.5 *μ*m) coupled to a Thermo Fisher Q-Exactive mass spectrometer. Mobile phase (A) consisted of 97% H_2_O, 3% ACN, 20 mM ammonium acetate, and 15 mM ammonium hydroxide pH 9.6. Organic phase (B) consisted of 100% ACN. Metabolites were resolved using the following linear gradient: 0 min, 85% B, 0.15 ml/min; 1.5 min, 85% B, 0.15 ml/min; 5.5 min, 40% B, 0.15 ml/min; 10 min, 40% B, 0.15 ml/min; 10.5 min, 40% B, 0.3 ml/min; 14.5 min, 40% B, 0.3 ml/min; 15 min, 85% B, 0.15 ml/min; 20 min, 85% B, 0.15 ml/min. The mass spectrometer was operated in positive ionization mode with a MS1 scan at resolution = 70,000, automatic gain control target = 3 x 10_6_, and scan range = 60-186 m/z and 187-900 m/z. This protocol was adapted from Mentch et al., 2015. Individual metabolite data were called using MAVEN (Melamud et al., 2010; Clasquin et al., 2012) with retention times empirically determined in-house. Peak Area Top values were analyzed to determine metabolite expression.

#### LC-MS DNA Methylation Assay

500 ng of cytosine, 5mC, and 5hmC double-stranded DNA standards from Zymo Research were incubated with Zymo Research DNA Degradase Plus for 3 hours at 37°C following enzyme inactivation at 70°C for 20 minutes. 75 *μ*l of ACN was added to each standard reaction to bring the final volume to 100 *μ*l. DNA standards were centrifugated at 21,100xg for 5 minutes at 4°C after which 75 *μ*l of supernatant was transferred to a glass vial for LC-MS analysis. A range of 0.1 to 100 ng of digested DNA was analyzed via LC-MS to determine each nucleoside’s linear range of detection.

Genomic DNA extracted using a Promega Wizard Genomic DNA Purification Kit after which 1 *μ*g was incubated with Zymo Research DNA Degradase Plus at 37°C following enzyme inactivation at 70°C for 20 minutes. 175 *μ*l of ACN was added to each reaction to bring the final volume to 200 *μ*l. DNA samples were centrifugated at 21,100xg for 5 minutes at 4°C after which 150 *μ*l of supernatant was transferred to a glass vial for LC-MS analysis.

Each prepared nucleoside sample was injected onto a Thermo Fisher Scientific Vanquish UHPLC with a Waters XBridge BEH Amide column (100 m x 2.1 mm, 3.5 *μ*m) coupled to a Thermo Fisher Q-Exactive mass spectrometer run in positive ionization mode. Mobile phase (B) consisted of H_2_O, 3% ACN, 20 mM ammonium acetate, and 15 mM ammonium hydroxide pH 9.6. Organic phase (B) consisted of 100% ACN. Nucleosides were resolved using the following linear gradient: 0 min, 85% B, 0.15 ml/min; 1.5 min, 85% B, 0.15 ml/min; 5.5 min, 40% B, 0.15 ml/min; 10 min, 40% B, 0.15 ml/min; 10.5 min, 40% B, 0.3 ml/min; 14.5 min, 40% B, 0.3 ml/min; 15 min, 85% B, 0.15 ml/min; 20 min, 85% B, 0.15 ml/min. The mass spectrometer was operated in positive ionization mode with a MS1 scan at resolution = 70,000, automatic gain control target = 3 x 10_6_, and scan range = 200-300 m/z, followed by a DDA scan with a loop count of 5. DDA settings were as follows: window size = 2.0 m/z, resolution = 17,500, automatic gain control target = 1 x 10_5_, DDA maximum fill time = 50 ms, and normalized collision energy = 30. This protocol was adapted from Mentch et al., 2015. Individual metabolite data were called using MAVEN (Melamud et al., 2010; Clasquin et al., 2012) with retention times empirically determined in-house. Peak Area Top values were analyzed to determine metabolite expression with dAdenosine being utilized for internal normalization (m/z dCytosine: 228.0978, m/z d5mC: 242.1135, m/z d5hmC: 258.108, m/z dAdenosine: 252.109).

#### Histone Isolation and Chemical Derivatization

Tissue culture cell pellets were resuspended in 800 *μ*l of ice-cold Buffer A (10 mM Tris-HCl pH 7.4, 10 mM NaCl, and 3 mM MgCl_2_) supplied with protease and histone deacetylase inhibitors (10 *μ*g/ml leupeptin, 10 *μ*g/ml aprotinin, 100 *μ*M phenylmethylsulfonyl fluoride, 10 mM nicotinamide, 1 mM sodium-butyrate, and 4 *μ*M trichostatin A) followed by 80 strokes of light pestle homogenization in a 1 ml Wheaton dounce homogenizer. Cell homogenate was then transferred to a 1.5 ml microcentrifuge Eppendorf tube and centrifugated at 800xg for 10 minutes at 4°C to pellet nuclei. The supernatant was either transferred to a fresh 1.5 ml Eppendorf tube or discarded. The nuclei pellet was resuspended in 500 *μ*l ice-cold PBS pH 7.4 followed by centrifugation at 800xg for 10 minutes at 4°C. The supernatant was discarded and nuclei were again washed with 500 *μ*l ice-cold PBS pH 7.4. Pelleted nuclei were then resuspended in 500 *μ*l of 0.4N H_2_SO_4_ and rotated at 4_o_C for 4 hours. Samples were centrifugated at 3,400xg for 5 minutes at 4°C to pellet nuclear debris and precipitated non-histone proteins. The supernatant was transferred to a new 1.5 ml Eppendorf tube after which 125 *μ*l of 100% trichloroacetic acid was added and incubated overnight on ice at 4°C. Samples were centrifugated at 3,400xg for 5 minutes at 4°C to pellet precipitated histone proteins. The supernatant was discarded, after which the precipitant was washed with 1 ml ice-cold acetone +0.1% HCl. Samples were centrifugated at 3,400xg for 2 minutes at 4°C and the supernatant was discarded. This process was repeated except precipitant was washed with 100% ice-cold acetone. Residual acetone was allowed to evaporate at room temperature for 10 minutes after which the dried precipitant was dissolved in 125 *μ*l H_2_O. Samples were then centrifugated at 21,100xg for 5 minutes at 4°C to pellet any remaining insoluble debris. Supernatant containing purified histone was then transferred to a new 1.5 ml Eppendorf tube and stored at −20°C until needed for future analyses.

For *M. musculus* liver tissue, an average tissue weight of 50 mg was homogenized in 800*μ*l ice-cold Buffer supplemented with protease and HDAC inhibitors (listed above) using a 1 ml Wheaton dounce homogenizer (20 strokes of the loose pestle followed by 20 strokes of the tight pestle) and strained through a 100 *μ*M filter before being transferred to a new 1.5 ml Eppendorf tube. All remaining steps for completing the histone isolation were performed as described previously in the tissue culture isolation protocol.

To prepare purified histone proteins for LC-MS/MS analysis, 5 *μ*g of histone was diluted with H_2_O to a final volume of 9 *μ*l. 1 *μ*l of 1 M triethylammonium bicarbonate was added to each sample to buffer the solution to a final pH of 7-9. Next, 1 *μ*l of 1:100 propionic anhydride:H_2_O was added to each sample and allowed to incubate for 2 minutes at room temperature. The propionylation reaction was then quenched via the addition of 1 *μ*l 80 mM hydroxylamine which was allowed to incubate for 20 minutes at room temperature. Next, propionylated histones were digested with 0.1 *μ*g trypsin for 4 hours at 37°C. Upon completion of trypsin digestion, 5 *μ*l of .02 M NaOH was added to bring the final pH between 9-10. Propionylated, trypsin digested histone peptides were then N-terminally modified with 1 *μ*l 1:50 phenyl isocyanate:ACN for 1-hour at 37°C. Modified peptides were desalted and eluted-off of Empore™C18 extraction membrane. Eluted samples were dried completely using a Thermo Fisher Scientific Savant ISS110 SpeedVac, resuspended in 40 *μ*l sample diluent (94.9% H_2_O, 5% ACN, .1% TFA), and transferred into glass vials for LC-MS/MS analysis.

#### Histone Proteomics Analysis

Derivatized histone peptides were injected onto a Dionex Ultimate3000 nanoflow HPLC with a Waters nanoEase UPLC C18 column (100 m x 150 mm, 3*μ*m) coupled to a Thermo Fisher Q-Exactive mass spectrometer at 700 nL/ min. Aqueous phase (A) consisted of H_2_O + 0.1% formic acid while the mobile phase (B) consisted of acetonitrile + 0.1% formic acid (B). Histone peptides were resolved using the following linear gradient: 0 min, 2.0% B; 5 min, 2.0% B; 65 min, 35% B; 67 min, 95% B; 77 min, 95% B; 79 min, 2.0% B; 89 min, 2.0% B. Data was acquired using data-independent acquisition (DIA) mode. The mass spectrometer was operated with a MS1 scan at resolution = 35,000, automatic gain control target = 1 x 10_6_, and scan range = 390-910 m/z, followed by a DIA scan with a loop count of 10. DIA settings were as follows: window size = 10 m/z, resolution = 17,500, automatic gain control target = 1 x 10_6_, DIA maximum fill time = AUTO, and normalized collision energy = 30.

DIA Thermo .raw files were imported into Skyline open source proteomics software (MacLean et al., 2010) and matched against a pre-constructed peptide database containing unique peptide proteoforms from H3 and H4. The identity of each individually called peak was verified by hand after which total MS1 peak area values were exported for downstream analyses. To control for differences in ionization efficiencies across histone peptides, all peptides possessing an identical primary sequence were analyzed as a peptide family which enabled the calculation of stoichiometric values for each unique peptide proteoform within the family. Stoichiometric values were compared across conditions to identify altered histone peptide abundances. Furthermore, combinatorial peptide stoichiometry values were deconvoluted by totaling the stoichiometry values for all peptides possessing a given PTM independent of any other PTMs that may be present on other residues of the same primary peptide. A Welch’s T-Test with a p-value cutoff of 0.05 was used to determine statistical significance.

Hierarchical clustering was performed in MATLAB R2016a using the ‘clustergram’ command. Pearson’s Correlation Coefficients were calculated in MATLAB R2016a using the ‘corr’ command.

#### RNAi Mediated Inhibition of MAT2A and H3K9 HMT Expression

For 35 mm tissue culture dishes, 1.5×10_5_ HCT116 cells were reverse transfected with 30 pmol of targeted or control RNAi using 5 *μ*l of Lipofectamine RNAiMAX in addition to 500 *μ*l Opti-MEM and 2.5 ml RPMI-1640 +10% FBS. MAT2A-RNAi treated HCT116 cells used for LC-MS metabolite analysis were cultured using dialyzed FBS.

#### Methionine Restriction of Tissue Culture Cells

All tissue culture cells were initially cultured in RPMI-1640 +10% FBS at 37°C, 5% CO_2_. One day before Met-restriction, 3.5×10_6_ cells were seeded into 100 mm tissue culture dishes. One-hour before the restriction began, RPMI-1640 was replaced with fresh, Met replete media. At the time of Met-restriction, cells were rinsed 2x with 3 ml of sterile PBS pH 7.4 followed by addition of RPMI-1640 void of Met. Any small molecule inhibitors or vehicle controls would be added at this time.

For experiments in which Met-restriction was followed by Met repletion, HCT116 cells were washed with PBS pH 7.4 2x to maximize small molecule inhibitor and DMSO removal before reintroduction of Met replete RPMI-1640.

HCT116 cells cultured for LC-MS metabolite analysis were cultured in RPMI-1640 containing 10% dialyzed FBS beginning 1-hour prior to beginning Met-restriction.

#### SILAC in Tissue Culture

197 ml of SILAC RPMI-1640 +10% DFBS void of L-Arg and L-Lys was supplemented with 50mg isotopically heavy 3C_6_ 15N_4_ L-Arg HCl and 9.43mg isotopically light L-Lys to obtain final concentrations of 1.149 mM and 0.219 mM, respectively. HCT116 cells were cultured in the SILAC media for 7 days (or 9 doubling cycles at a reported doubling time of 21 hours) after which cells were washed 3x with 5 ml PBS pH 7.4 before replacing the SILAC media with isotopically light L-Arg and L-Lys RPMI-1640 +10% DFBS void of Met. Half of the cells were treated with 5 *μ*M UNC0642 or 0.1% DMSO at this time. Furthermore, 0-hour control HCT116 cells which were never exposed to isotopically light L-Arg, Met-restricted RPMI-1640 were harvested. HCT116 cells cultured in the Met-restricted media were harvested after 24 hours.

#### SILAC Shotgun Proteomics Analysis

50 *μ*g of cytoplasmic protein extract from the previously described SILAC Met-restriction experiment were diluted with H_2_O into a final volume of 75 *μ*l. Next, dithiothreitol was added to each sample at a final concentration of 10 mM, followed by a 15-minute incubation at 95°C. A sonication water bath was used to resolubilize precipitated proteins post-heat denaturation. Iodoacetamide was then added to a final concentration of 20 mM, followed by a 20-minute incubation at room temperature protected from light. Alkylated proteins were precipitated with 337 *μ*l ice-cold acetone for 20 minutes at −20°C. Samples were centrifugated at 21,000xg for 15 minutes at 4°C, the supernatant was discarded, and precipitated protein was resuspended in 400 *μ*l of urea buffer (2 M urea and 100 mM ammonium bicarbonate pH 8). Precipitated proteins were again solubilized using a sonication water bath. Solubilized proteins were digested into peptides using 0.8 *μ*g trypsin at 37°C overnight. Upon digest completion, samples were desalted and eluted-off Empore™C18 extraction membrane. Eluted samples were dried completely using a Thermo Fisher Scientific Savant ISS110 SpeedVac, after which being resuspended in 50 *μ*l sample diluent (94.9% H_2_O, 5% ACN, .1% TFA) and transferred to glass vials for LC-MS/MS analysis.

Each prepared cytoplasmic peptide sample was injected onto a Dionex Ultimate3000 nanoflow HPLC with a Waters nanoEase UPLC C18 column (100 m x 150 mm, 3*μ*m) coupled to a Thermo Fisher Q-Exactive mass spectrometer at 800 nL/ min. Aqueous phase (A) consisted of H_2_O + 0.1% formic acid while the mobile phase (B) consisted of acetonitrile + 0.1% formic acid (B). Histone peptides were resolved using the following linear gradient: 0 min, 5.0% B; 5 min, 5.0% B; 120 min, 30% B; 122 min, 95% B; 127 min, 95% B; 129 min, 5.0% B; 145 min, 5.0% B. Data was acquired using data-dependent acquisition (DDA) mode. The mass spectrometer was operated with a MS1 scan at resolution = 70,000, automatic gain control target = 1 x 10_6_, and scan range = 400-1000 m/z, followed by a DDA scan with a loop count of 20. DDA settings were as follows: window size = 2.0 m/z, resolution = 17,500, automatic gain control target = 2 x 10_5_, DDA maximum fill time = 150 ms, and normalized collision energy = 28.

#### MaxQuant Proteomics Analysis

Thermo .raw DDA files were analyzed using MaxQuant v1.6.2.6 (Cox et al., 2014) using the default settings with the following exceptions: Labels: 1, Arg10; Instrument: Orbitrap; LFQ: LFQ min. ratio count: 2, LFQ min. number of neighbors: 3, LFQ average number of neighbors: 6; FASTQ File: Proteome: *H. Sapiens* Proteome ID UP000005640.

Identified proteins with a Q-value of 0.01 or above were removed. For the remaining proteins, LFQ light (L-Arg_+0_) and heavy (L-Arg_+10_) values were summed to generate LFQ total protein abundance values. LFQ light values were divided by the corresponding LFQ total value to generate a percent abundance of LFQ light species for each identified protein. Welch’s t-Tests with a p-value cut off of 0.05 were used to determine significant accumulation of LFQ light species in Met-restricted HCT116 samples relative to 0-hour controls. For proteins determined to have significant LFQ light species abundance after Met-restriction, LFQ total values were used to determine the corresponding change in overall protein abundance. Up to 2 null-values were removed from replicates of an experimental group to account for missed identifications. If 3 of the 4 replicates for a single experimental group contained null-values, that protein was removed from all experiment groups and therefore excluded from any downstream analyses. A Welch’s t-Test with a p-value cut off of 0.05 was used to determine if an identified protein was significantly more abundant after Met-restriction relative to 0-hour controls.

#### SILAC Histone Proteomics Analysis

Histones were isolated, chemically derivatized, and analyzed by LC-MS/MS as described in “Histone Isolation and Chemical Derivatization” and “Histone Proteomics Analysis.” DDA Thermo .raw files of histone proteins with incorporated isotopically heavy 3C_6_ 15N_4_ L-Arg were used to construct an H3K9/K14 peptide library using Skyline opensource proteomics software (MacLean et al., 2010). DIA files were then matched against both isotopically heavy and light L-Arg peptide libraries after which peptide identities were confirmed by hand. MS1 intensity values for both isotopically heavy and light L-Arg peptides were summed to generate total MS1 peptide proteoform intensity values which were then used to calculate the percent of total MS1 peptide proteoform signal which was contributed by isotopically heavy peptide proteoforms. Welch’s T-Tests with a p-value cutoff of 0.05 were used to determine significant differences in total MS1 peptide proteoform stoichiometric values as well as the percent contribution of isotopically heavy peptide proteoforms to total peptide proteoform intensity values.

#### Chromatin Immunoprecipitation and Sequencing

30×10_6_ HCT116 cells were resuspended in 1 ml MNase digestion buffer (50 mM Tris-HCl pH 7.6, 1 mM CaCl_2_, and 0.2% Triton X-100) after which 330 *μ*l of the cell suspension was aliquoted into individual 1.5 ml tubes. Each tube was treated with either 45, 52.5, or 60 units of Worthington MNase for 5 minutes at 37°C shaking at 400 rpm to generate mono-nucleosomes. MNase digestions were stopped via the addition of EDTA to a final concentration of 5mM. Digested cells were then pooled and lysed at 4°C using a Covaris S220 Focused-ultrasonicator with the following settings: Peak Power = 75, Duty Factor = 5, Cycles/Burst = 100, Duration = 30” on/30” off x2. Insoluble debris was centrifugated at 10,000 rpm for 10 minutes at 4°C after which the supernatant containing prepared nucleosomes was transferred to a new tube. Glycerol was added to a final concentration of 5% prior to −80°C storage. For input controls, 43 *μ*l of prepared nucleosomes were aliquoted and saved for further processing at −80°C.

Next, 860 *μ*l of prepared nucleosomes was incubated with 5 *μ*g of the desired antibody, rotating overnight at 4°C. The nucleosome/antibody solution was added to 50 *μ*l of prepared M-280 Sheep Anti-Rabbit IgG superparamagnetic Dynabeads™ and allowed to rotate at 4°C for 4 hours. Conjugated beads were then washed for 10 minutes, rotating at 4°C under the following conditions: 3x 1 ml RIPA (10 mM Tris pH 7.6, 1 mM EDTA, 0.1% SDS, 0.1% Na-deoxycholate, 1% Triton X-100) buffer, 2x 1 ml RIPA buffer +0.3 M NaCl, 2x 1 ml LiCl buffer (0.25 M LiCl, 0.5% NP-40, 0.5% Na-deoxycholate), 1x with 1 ml 1xTE +50 mM NaCl. Following the final wash, beads were centrifugated at 4°C for 3 minutes at 960xg after which the supernatant was carefully removed and discarded.

To elute immunoprecipitated nucleosome species, beads were resuspended in 210 *μ*l of elution buffer (50 mM Tris-HCl pH 8.0, 10 mM EDTA, 1% SDS) and incubated at 65°C for 30 minutes shaking at 400 rpm. Beads were resuspended every 2 minutes via brief vortexing. Next samples were centrifugated at 16,000xg for 1 minute at room temperature after which 200 *μ*l of supernatant was transferred to a new 1.5 ml Eppendorf tube. Eluted nucleosome species were then treated with 2 *μ*l Thermo Fisher Scientific Proteinase K for 2 hours at 55°C. DNA was recovered from the Proteinase K digestion using the QIAquick PCR Purification Kit.

DNA libraries were prepared by the University of Wisconsin-Madison Biotechnology Center using the Illumina TruSeq DNA Nano kit and then sequenced across 4 lanes of an Illumina 1×100 HiSeq2500 High Throughput flowcell.

FASTQ files were aligned unpaired using bowtie2-2.1.0 (Langmead and Salzberg, 2012) against human genome 19. Bowtie2 output SAM files were converted to BAM files, sorted and indexed using samtools 1.5 (Li et al., 2009). Broad peaks were called from sorted BAM files using MACS2-2.1.1.20160309 (Zhang et al., 2008) with a Q-value cutoff of 0.1. Input DNA was used as the control file. All identified peaks possessing a log_2_ fold-change below 1.75 were discarded prior to downstream analyses. Bedtools v2.25.0 (Quinlan and Hall, 2010) was used to identify unique and overlapping across samples and also used to assign raw mapped reads to pre-specified genomic loci. Genomic loci were annotated using HOMERv4.9 (Heinz et al., 2010).

#### Quantitative PCR

RNA was extracted from HCT116 cells using Thermo Fisher Scientific TRIzol and from liver tissue using Sigma TRI Reagent, both by following the manufacturer’s instructions. The concentration and purity of RNA were determined by absorbance at 260/280 nm.

cDNA was prepared from 1 *μ*g of RNA using RevertAid Reverse Transcriptase for HCT116 RNA and Superscipt III for liver RNA. Oligo dT primers were used in both cDNA synthesizing reactions.

HCT116 cDNA was analyzed on a Roche 480 Light Cycler System with PerfeCTa SYBR Green SuperMix while cDNA generated from liver was analyzed on an Applied Biosystems StepOne Plus instrument with Sybr Green PCR Master Mix. HCT116 cDNA qPCR reactions were normalized internally using GAPDH while liver cDNA qPCR reactions were normalized internally using actin. Primer sequences are available in Table S3.

#### RNA sequencing

RNA libraries were prepared by Novogene using NEBNext adaptors followed by paired end 2×150 sequencing on their Illumina Platform. FASTQ files were trimmed using Trimmomatic v0.39 (Bolger et al., 2014) and aligned using RSEM v1.2.4 (Li et al., 2011) against the human genome 18 reference transcriptome. RSEM .genes output files were used for Ebseq EBMultiTest multiple comparison analysis (Leng et al., 2013). Gene lists associated with individual EBMultiTest patterns were further analyzed with GSEA 2 (Tamayo et al., 2005; Mootha et al., 2003)

#### MNase Accessibility Assay

6×10_6_ HCT116 cells were resuspended in 1 ml of digestion buffer (50 mM Tris-HCl pH 7.6, 1 mM CaCl_2_, and 0.2% Triton X-100) in addition to 15 unites of Worthington MNase. MNase treated cell suspensions were incubated for 5 minutes at 37°C shaking at 400 rpm. MNase digestions were stopped via the addition of EDTA to a final concentration of 5 mM. Digested cells were then pooled and lysed at 4°C using a Covaris® S220 Focused-ultrasonicator with the following settings: Peak Power = 75, Duty Factor = 5, Cycles/Burst = 100, Duration = 30” on/30” off x2. Insoluble debris was pelleted via centrifugation at 10,000 rpm for 10 minutes at 4°C after which the supernatant was transferred to a new tube. 150 *μ*l of digested chromatin was treated with 2 *μ*l of Thermo Fisher Scientific Proteinase K for 90 minutes at 55°C. DNA was then purified using the QIAquick PCR Purification Kit. Next, 1 *μ*g of purified DNA was separated on a 2% agarose gel and quantified using GelAnalyzer 2010a.

#### Western-Blotting and Image Quantification

Protein separated on SDS-PAGE gels were transferred onto 0.2 *μ*m nitrocellulose membrane and stained with 5 ml of LI-COR REVERT Total Protein Stain for 5 minutes at room temperature before being blocked in 0.1% fat-free milk PBS pH 7.4 for 1 hour. Primary antibodies were diluted 1:1000 in 5% fat-free milk PBST pH 7.4 and incubated with the membrane overnight at 4°C. LI-COR secondary antibodies were used at a 1:15,000 dilution in 5% fat-free milk PBST pH 7.4 and incubated with the membrane for 1 hour at room temperature. Membranes were imaged using an Odyssey Infrared Imager (model no. 9120). Densitometry was performed using Image Studio™ Lite software. All data was normalized to REVERT™ Total Protein Stain.

#### In vivo Tests

For glucose tolerance tests, mice were fasted overnight for 16 hours and then injected with glucose (1 g/kg) intraperitoneally as previously described (Arriola Apelo et al., 2016). Glucose measurements were taken using a Bayer Contour blood glucose meter and test strips. Mouse body composition was determined using an EchoMRI Body Composition Analyzer (EchoMRI_TM_, Houston, TX, USA) according to the manufacturer’s procedures.

#### Protein Purification

The *H. sapiens* Ehmt1 catalytic subunit *E. coli* expression plasmid used in this study (Addgene plasmid # 51314) was acquired from addgene, made possible through a gift by Cheryl Arrowsmith. The *H. sapiens* Ehmt2 catalytic subunit *E. coli* expression plasmid used in this study was provided by the laboratory of Peter W. Lewis at the University of Wisconsin-Madison.

Transformed Rosetta™ *E. coli* competent cells were cultured in 1L of 2XYT media at 37°C to an O.D. 600 of 0.8. IPTG was then added to each culture at a final concentration of 1mM. Cultures were allowed to grow for 16 hours at 18°C before being harvested and stored at −80°C. Pellets were resuspended in 30ml Buffer A (50 mM NaPi, 250 mM NaCl, 10mM imidazole, pH 7.2) and lysed using sonicated in the presence of lysozyme. Lysate was centrifugated and the supernatant was collected. The supernatant was then loaded onto a HisTrap FF nickel column in line with a GE AKTA FPLC. Protein was eluted off the column using a linear gradient ending in 100% Buffer B (50 mM NaPi, 250 mM NaCl, 250 mM imidazole, pH 7.2) and collected into time-dependent fractions. Fractions were analyzed by coomassie staining of SDS-PAGE gels to assess purity.

#### Histone Methyltransferase Activity Assay

The methyltransferase activity of recombinantly purified Ehmt1 and Ehmt2 catalytic subunits was assessed using Promega Corporation’s MTase-Glo™ Methyltransferase Assay. Assays were performed using 12.5 nM enzyme in the presence of saturating concentrations of SAM (30 *μ*M) and unmodified H3K9 peptide (30 *μ*M, WARTKQTARKSTGGKAPR – 3’) as well as either saturating levels of UNC0642 (5 *μ*M) or 0.1% DMSO. Reactions proceeded for 1 hour at 30°C before addition of MTase-Glo™ reagent following the Promega protocol. Luminescence was detected using a Biotek Synergy H4 Hybrid plate reader. All values detected were determined to be within the instrument’s linear range of detection via the use of a SAH standard curve. To test for reversibility of enzyme inhibition by small molecule, 1 *μ*M Ehmt1 and Ehmt2 were incubated with saturating concentrations of UNC0642 (5 *μ*M) or 0.1% DMSO followed overnight 1,000x or 10,000x dialysis with a molecular weight cutoff of 10,000 Da against 1x Reaction Buffer without BSA (20mM Tris, pH 8.0, 50mM NaCl, 1mM EDTA, 3mM MgCl_2_, 1mM DTT). Following dialysis, enzyme and reaction components were diluted using the appropriate dialysis buffers to the working concentrations described above. The MTase-Glo™ assay was then repeated using identical conditions as described above.

#### Quantification and Statistical Analysis

Explanations of statistical analyses and parameters are present in the main and supplemental figure legends as well as in ‘Method Details” where appropriate.

#### Data and Software Availability

RNA-sequencing as well as H3K9me1 and H3K9me3 ChIP-sequencing data have been uploaded to Gene Expression Omnibus (GEO) and will be made publicly available upon manuscript publication.

## KEY RESOURCES TABLE

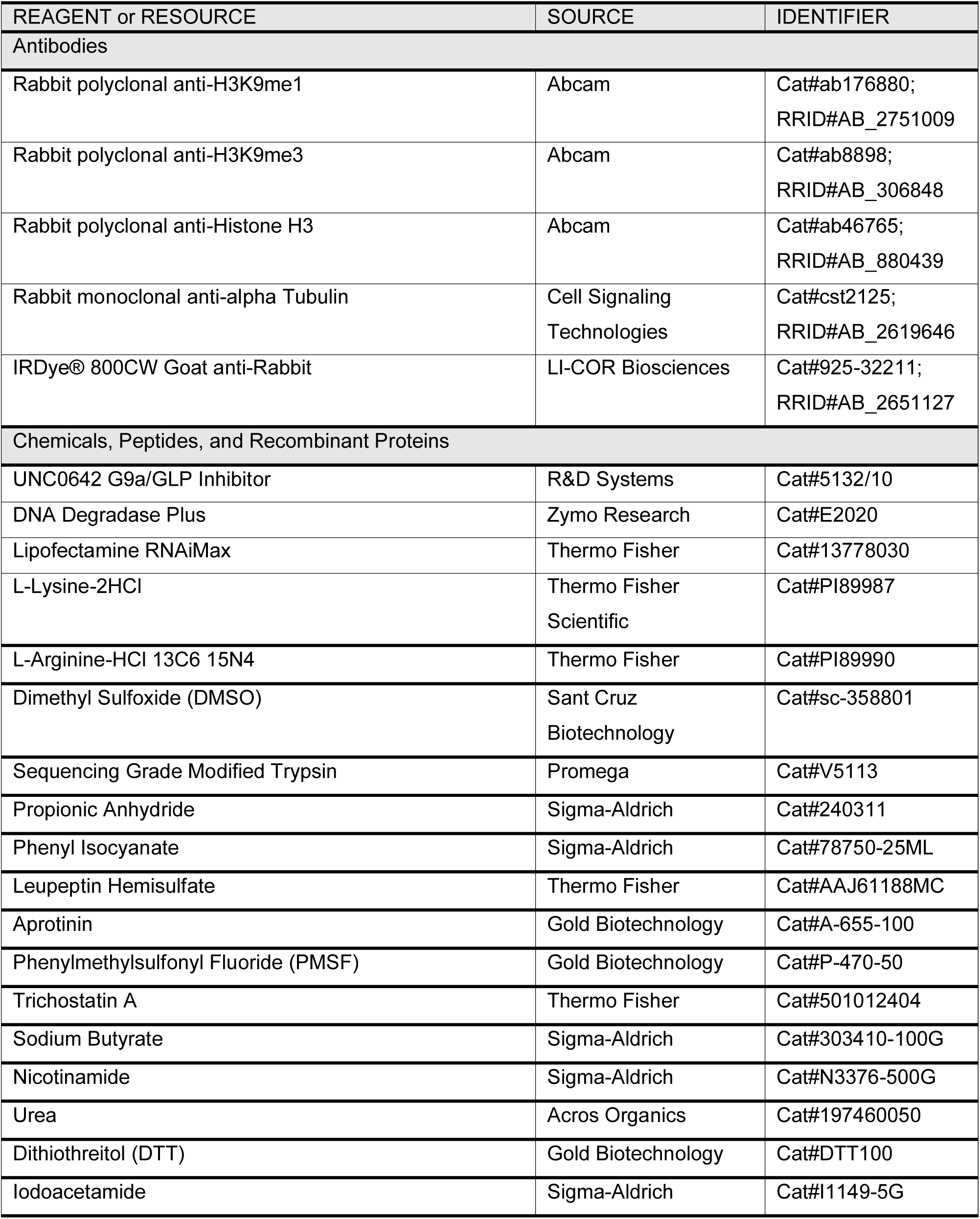
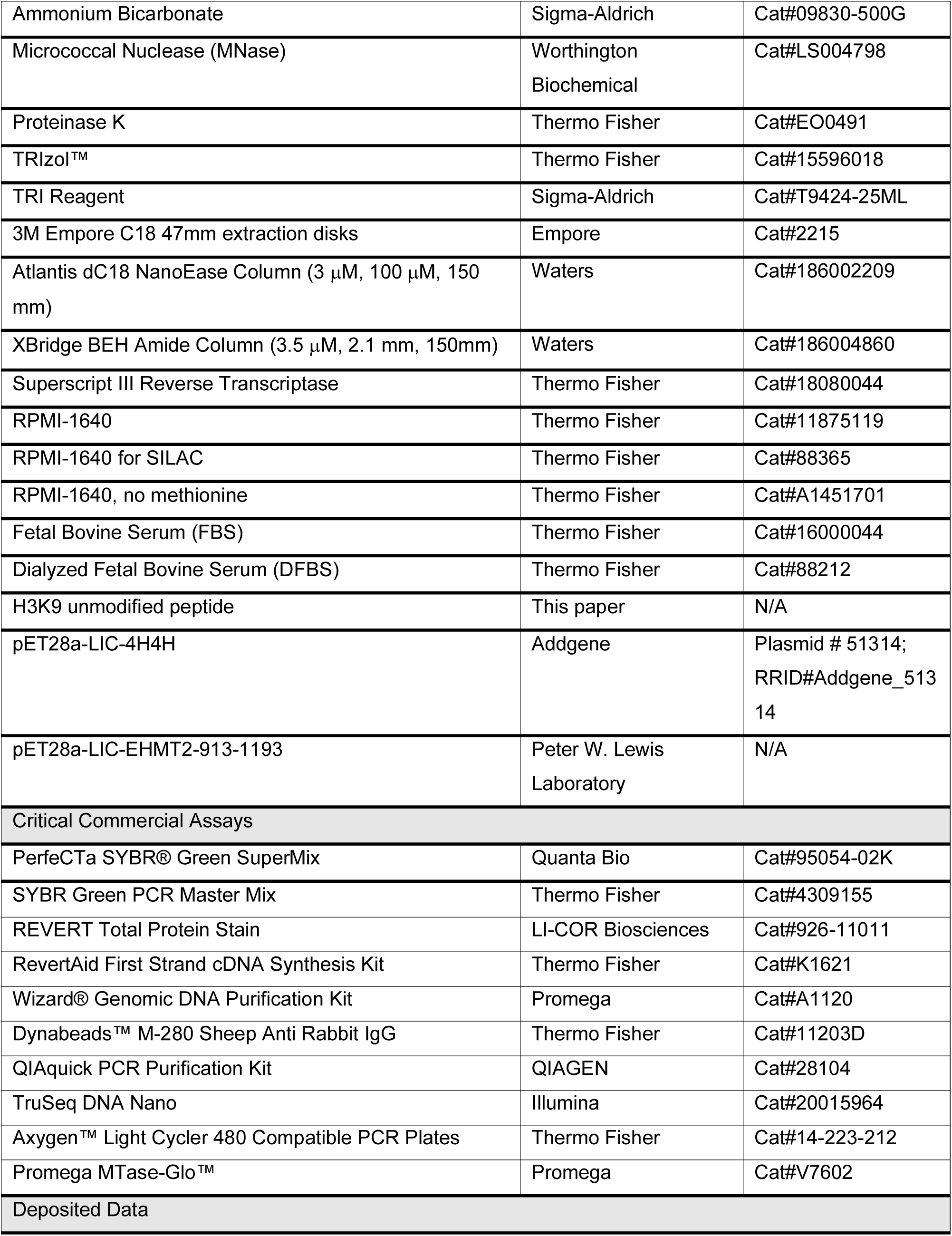
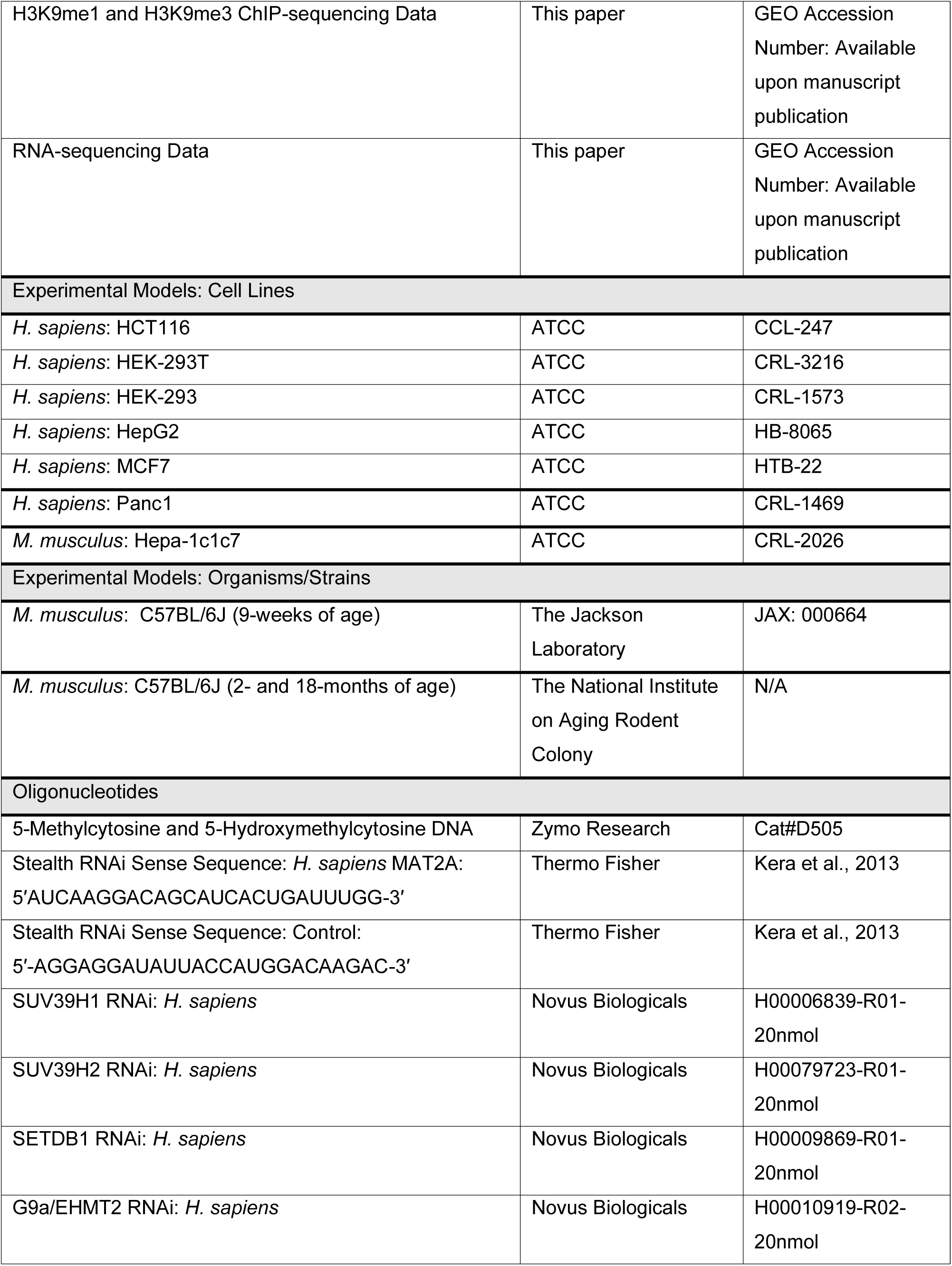
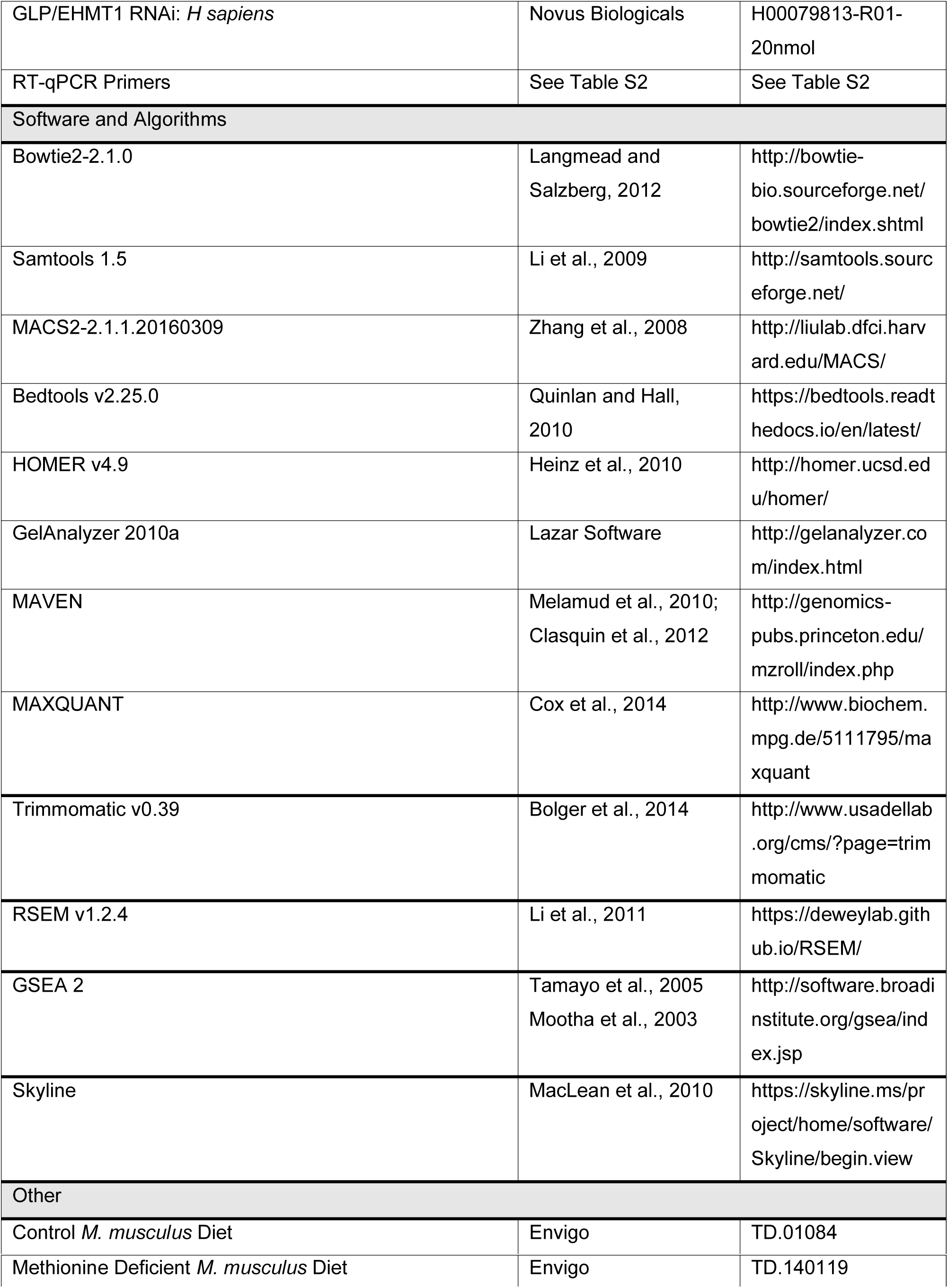

## Supplemental File Titles and Legends

Document S1, Related to Figures 3 and S3. Primary Antibody and REVERT Total Protein Stain Western Blot Membranes.

*(A)* Cytoplasm anti-H3K9me1 (Figure 3E). (*B*) REVERT Total Protein Stain for cytoplasm anti-H3K9me1 (Figure 3E). (*C*) Cytoplasm anti-H3 (Figure 3E). (*D*) REVERT Total Protein Stain for cytoplasm anti-H3 (Figure 3E). (*E*) Cytoplasm anti-*α*Tubulin (Figure 3E). (*F*) REVERT Total Protein Stain for cytoplasm anti-*α*Tubulin (Figure 3E). (*G*) Nucleoplasm anti-*α*Tubulin (Figure 3E). (*H*) REVERT Total Protein Stain for nucleoplasm anti-*α*Tubulin (Figure 3E). (*I*) Purified histone anti-H3K9me1 (Figure S3F). (*J*) REVERT Total Protein Stain for purified histone anti-H3K9me1 (Figure S3F).

Table S1, Related to Figures 1-3, 6-7, S1-S3, and S5. LC-MS/MS % Abundance Histone PTM Data.

Table S2, Related to Figures 2, 5, 7, S3, and S5. RT-qPCR Primers.

Table S3, Related to Figure 3. SILAC Cytoplasmic Proteins.

