## Supplementary figures and images for "Methyl-Metabolite Depletion Elicits Adaptive Responses to Support Heterochromatin Stability and Epigenetic Persistence"

### Document S1

Document S1. Primary Antibody and REVERT Total Protein Stain Western Blot Membranes.

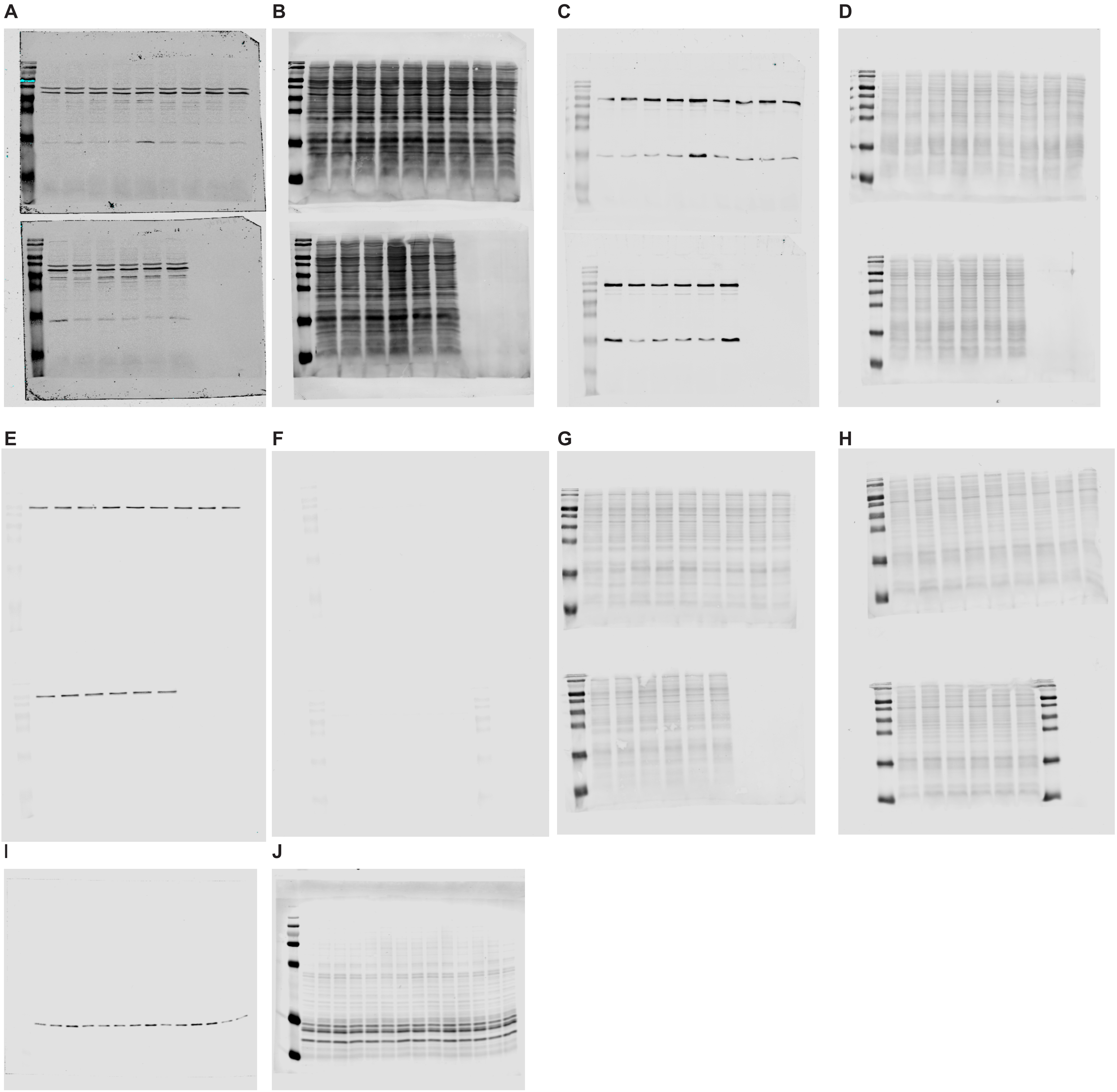
